# Combined interventions are required to reverse biodiversity and carbon losses in tropical forest frontiers

**DOI:** 10.1101/2025.11.04.686271

**Authors:** Leonardo S. Miranda, James R. Thomson, Joice Ferreira, Toby A. Gardner, Erika Berenguer, Alexander C. Lees, Ralph Mac Nally, Luiz E. O. C. Aragão, Pedro H. S. Brancalion, Silvio F. B. Ferraz, Rachael D. Garrett, Paulo G. Molin, Nárgila G. Moura, Sâmia S. Nunes, Luke Parry, Juliana M. Silveira, Ima C. G. Vieira, Cecilia Viana, Jos Barlow

## Abstract

Halting deforestation and promoting restoration are at the core of biodiversity and climate strategies in tropical forest regions. Avoiding forest disturbances is also critically important but has received far less attention, and there is a lack of clarity about the cost-effectiveness of these three different interventions. We provide the first comparison of the biodiversity and carbon benefits and costs associated with each intervention, comparing observed and counterfactual outcomes based on in-depth field assessments and high-resolution remote sensing in Amazonian Brazil. Avoiding forest disturbances delivered the greatest benefits and was more cost-effective than either avoiding deforestation or restoration, with results being robust to a range of benefit and cost assumptions. However, combined interventions delivered the greatest gains and were essential to reverse biodiversity and carbon losses.

---

Tropical primary forests have a disproportionate role in conserving global biodiversity, mitigating climate change and supporting human well-being *(1,2,3)*. However, rates of tropical forest loss remain persistently high despite a proliferation of policies and investments to conserve them in recent decades *(4,5)*. Remaining forests, including those within existing protected areas, are often threatened by widespread degradation resulting from human-driven disturbances, including forest fires exacerbated by climate change, logging and edge effects *(6,7)*. Deforestation and disturbances are driving widespread biodiversity loss and turning tropical forest regions from carbon sinks to net sources *(8,9)*.

Halting and reversing this decline requires actions across the entire mitigation hierarchy to avoid further: (i) deforestation and (ii) human disturbances, and (iii) restore already deforested areas *(10,11)*. Preventing deforestation is the most preeminent forest conservation intervention, promoted through global programs, such as the REDD+ mechanism under the United Nations Framework Convention on Climate Change (UNFCCC; *12)*, national legislation, such as Brazil’s National Vegetation Protection Law (NVPL; *13)* and the Action Plan for the Prevention and Control of Deforestation in the Brazilian Amazon (PPCDAm; *14)*, and due diligence requirements in commodity trade, such as the European Union Deforestation Regulation (EUDR; *15)* and private sector voluntary zero-deforestation commitments *(16,17)*. Interest in restoration has grown rapidly, reflected in the United Nations Decade on Ecosystem Restoration *(18)* and the Bonn Challenge *(19)*, expanding biodiversity and carbon markets, and government programs, such as the NVPL and the Brazil’s Plan for Native Vegetation Recovery (Planaveg; 12 million ha goal; *20)*. Although the importance of avoiding forest disturbances was formally recognized in climate-mitigation strategies by the UNFCCC in 2007, it has attracted far less attention and investment than initiatives focused on preventing deforestation or restoring cleared land *(12,16)*, held back by its perceived lesser importance, difficulties in monitoring at scale, challenges in assigning responsibility for forest disturbances such as fires, and the difficulties of demonstrating additionality from interventions targeted at avoiding disturbances per se (6,*21-25)*. These challenges are exacerbated by the broad range of disturbance types that result in forest degradation; while landscape-scale drivers of degradation such as edge effects and fragmentation (size and isolation) are tightly coupled with deforestation, within-forest disturbances such as fire and logging, can occur independently of forest loss *(8,25)*.

Despite these challenges, the growing prevalence and severity of forest disturbances demonstrates a clear and increasingly urgent need to include countermeasures within the forest conservation toolbox *(7,21,25)*. However, this need is currently obfuscated by the lack of information on the environmental benefits and economic costs of avoided disturbance compared to avoided deforestation or restoration. Furthermore, it is not clear to what extent single or combined approaches are needed to bring about net-positive outcomes for biodiversity and carbon. Here, we address these gaps using in-depth field data and a novel analytical framework that estimates the landscape-wide benefits and costs of interventions aimed at preventing and reversing biodiversity loss and mitigating climate change. While this question is relevant for tropical forests globally, we focus our analysis on the consolidated deforestation frontier of the Brazilian Amazon where the trade-offs over different types of intervention are made most acute due to continued forest loss, extensive forest fires and logging activities, and the availability of deforested land for restoration.

We quantified the biodiversity and carbon benefits that could be derived from the successful implementation of three major conservation interventions: *avoid deforestation, avoid within-forest disturbances* (hereafter *avoid forest disturbances)*, and *restoration* to meet legal compliance (Table 1). Using a counterfactual approach, they were evaluated individually and in combination, for all forests (primary and secondary) and for primary forests, for two ecologically distinct regions within the Arc of Deforestation: Santarém and Paragominas (Supplementary Material section 1.1, fig. S1, *26)*. Together these cover over 3 million ha and capture broad gradients of deforestation history, forest condition, land-use pressures, and legal compliance *(26-29)*.

**Table 1.** Conceptual framework. How each counterfactual intervention scenario relates to 2010 baseline conditions, 2020 landscape observed changes, the landscape effects, expected biodiversity and carbon benefits, and the relevant legal frameworks.

| **Intervention**<br>**(Counterfactual**<br>**scenario)** | **2010 Landscape**<br>**(Baseline)** | **2020 Landscape**<br>**(Observed Change)** | **Landscape effects if**<br>**intervention applied** | **Legal Frameworks &**<br>**Policy Instruments** |
| --- | --- | --- | --- | --- |
| Avoid Deforestation | Regions had a mix of undisturbed and disturbed primary forests, secondary forests, and deforested land; areas under pressure from agricultural expansion. | 159,000 ha deforested during this period (18,709 ha in secondary forests), mainly in areas already affected by disturbance (e.g., fire, logging, and edge effects). | All deforestation from 2010 onward is avoided; no direct habitat loss and new edge effects. | Brazilian Native Vegetation Protection Law (NVPL); Action Plan for Deforestation Prevention (PPCDAm); national and subnational land-use licensing of forest use. |
| Avoid forest Disturbances | Widespread presence of forest fires and evidence of logging in primary forests; governance gaps in managing disturbances. | 273,600 ha disturbed forests by fire or logging in 10 years; 66.5% of regenerating secondary forests were disturbed by fire. | All within-forest disturbances (i.e., fire and logging) were avoided; improvements in forest condition. | Federal and state legislation to prevent illegal fires (Law 12651/2012; Law 9605/1998); fire control policies (Decree 2661/1998); and forest management regulations (Law 9605/1998; IN 05/2006). |
| Restoration to Legal Compliance | 18% of private properties had forest deficits relative to legal requirements; absence of structured restoration plans; no guidance promoting natural regeneration near forest remnants. | 80,705 ha of new forests are required to resolve legal restoration deficit, according to the multiple criteria in NVPL. | Secondary forests are allocated to meet NVPL thresholds, starting with APPs then RLs; prioritize proximity to existing forests to reduce edge effects and promote regrowth. | NVPL requirements for APP and RL restoration; Planaveg, Low Carbon Agriculture Plan; REDD+ initiatives; international targets (e.g., Bonn Challenge, GBF Target 2). |

First, we mapped the observed land-use transitions between 2010 and 2020 across the study regions. Analysis of 30-m resolution land cover data revealed 158,980 ha of deforestation (63,787 ha of secondary forests), 273,558 ha of primary forest fire or logging, and 129,505 ha of secondary forests regrowth (63.1% of which were subsequently disturbed by fire) (Supplementary Material sections 1.2 and 1.3, figs. S1-S6, *26)*. These changes formed the baseline scenario of no additional interventions and provided the basis of our *avoid deforestation* and *avoid forest disturbance* scenarios, which prevented all deforestation or all fire and fire and logging, respectively (Supplementary Material section 1.4, *26)*. Landscape-scale disturbances such as edge effects were determined by landscape configuration and are therefore present in all scenarios. We also developed a property-level *restoration* scenario, using a novel approach to ensure all 6,900 registered rural properties in the regions meet their legal compliance of forest cover (depending on multiple criteria under the NVPL; *13)*. We first restored mandatory Permanent Protection Areas (APPs, acronym in Portuguese), then allocated Legal Reserve (RLs) deficits within properties prioritizing areas near existing forests to enhance natural regeneration and reduce edge effects, adding 80,705 ha of secondary forests compared to the baseline (Supplementary Material section 1.5, *26)*.

To measure the biodiversity and carbon outcomes of each scenario, we used random forest models trained on extensive field data collected in 2010, covering 381 0.25 ha transects grouped in 38 catchments (Supplementary Material section 1.6, *26)*. Occurrence probabilities (160 bird and 426 tree species) and above-ground carbon were modelled as functions of forest type (primary, disturbed, secondary), forest condition (age, time since last disturbance, edge density), and landscape context (distances to edge, road and river, climate, elevation) (table S1, fig. S7). Models were tested using a k-fold cross-validation scheme in which catchments were iteratively held out of training. Performance was assessed using the area under receiver operator curves (AUC), for biodiversity; and root mean square error (RMSE) for carbon (AUC > 0.8 for most species [table S2], RMSE 43.8 Mg C ha−^1^ for carbon). These models produced pixel-level predictions of occurrence probabilities and carbon stocks for the 2010 landscape (baseline), for the 2020 landscape (observed change), and for each intervention scenario (figs. S8-S12). For biodiversity, a synthetic value was calculated as the average weighted occurrence probability for all species, with weights inversely proportional to each species’ geographic range size to emphasize the conservation value of range-restricted taxa (Supplementary Material section 1.7, *26)*. Net changes in biodiversity and carbon were calculated as the difference between the summed values of all pixels in the 2010 baseline and the pixels in 2020 under each scenario. The benefits of each intervention were then calculated as the difference between the counterfactual intervention scenario and observed change in 2020. Although the immediate and successful implementation of each intervention is highly unlikely to occur over short timescales in the real world, we intentionally use these idealized outcomes to compare the maximum and per-ha benefits.

For both regions, all interventions helped to avoid or partially compensated for the predicted losses of biodiversity and carbon (Fig. 1, fig. S13). Avoiding either deforestation or forest disturbances delivered the greatest benefits, with avoided deforestation mitigating 28.0% and 60.8% of biodiversity and carbon loss, respectively, and avoided forest disturbances mitigating 68.5% and 46.8% of biodiversity and carbon loss.

**Fig 1.**
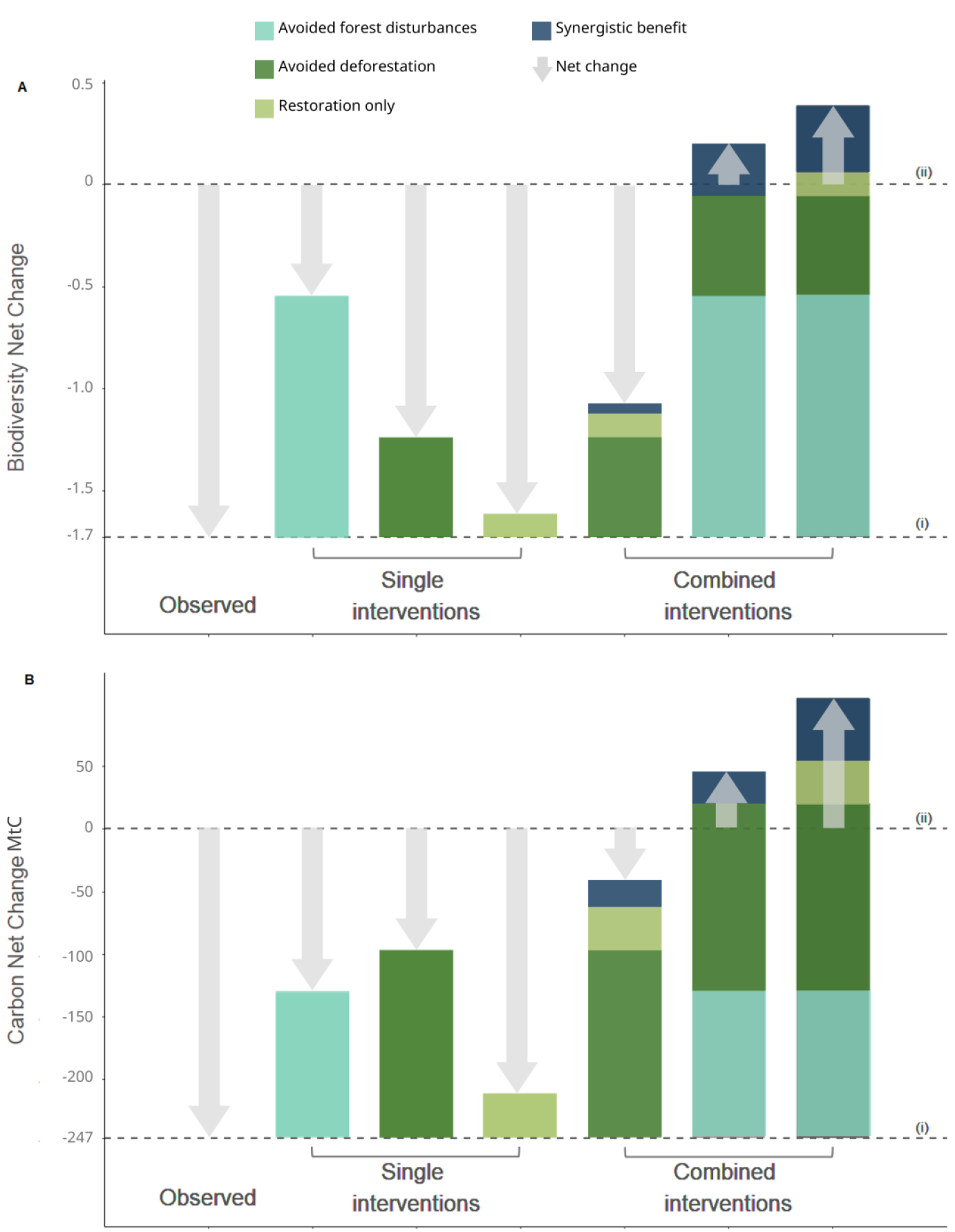
Summed net change and benefits for biodiversity and carbon under different conservation intervention scenarios. Panels represent the total net change and benefits for biodiversity (A) and carbon (B) over a 10-year period (2010–2020). Y-axis show the summed average weighted occurrence probabilities and summed megagrams of carbon, respectively. Gray arrows indicate the net change for each scenario, with downward arrows denoting net losses and upward arrows indicating net gains. Avoided degradation (light blue bar), avoided deforestation (green bar), and restoration (light green bar) all contribute to benefits, with the combined interventions generating synergistic gains (dark blue), which exceed the sum of individual interventions. Dashed lines represent the 2010 baseline (upper line) and the predicted changes from 2010 to 2020 (lower line).

Restoration delivered the lowest summed benefits and, when implemented without avoiding deforestation or forest disturbances, only compensated for 6.5% and 14.7% of the observed loss of biodiversity and carbon, respectively. Crucially, net-positive, landscape-wide changes were only achieved by combining both avoided forest disturbances and avoided deforestation (Fig. 1), enhancing biodiversity and carbon by 12% and 18% above the 2010 baseline, respectively. Implementing all three interventions would have enhanced biodiversity and carbon by 22% and 42% above the baseline, respectively. The benefits accrued from combined interventions were not additive: between 15-19% for biodiversity and 10-20% of the total combined benefits for carbon resulted from synergies between different interventions (Fig. 1).

The benefits of each intervention were highly variable, with pixel-level change varying according to the location and metric assessed (Fig. 2, fig. S14). For example, although avoided deforestation delivered the greatest summed benefits for carbon, it was less effective than avoiding forest disturbance for biodiversity. This likely reflects its predominant location in forests that had already been affected by edge effects (<500 m of forest edges; fig. S15A) or with a high prevalence of previous disturbance (fig. S15B). Biodiversity would be particularly sensitive to this as many forest species are sensitive to disturbances such as edge effects *(30)* or fire *(6)*.

**Fig. 2.**
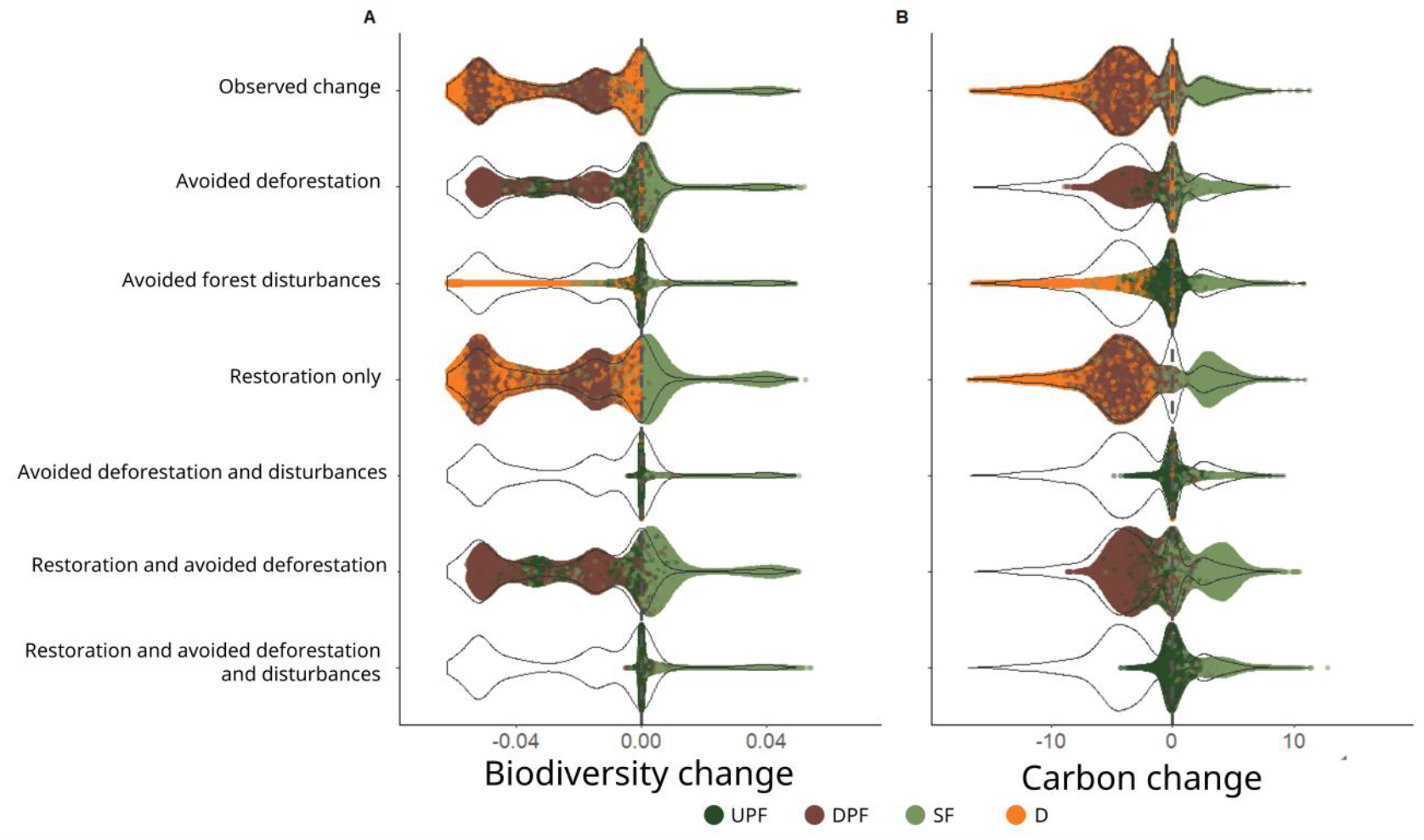
Pixel-level distribution of net changes in biodiversity and carbon across conservation intervention scenarios. Plots illustrate the pixel-level distribution of net changes in biodiversity (A) and carbon (B) with different scenarios. The y-axis displays the scenarios, while the x-axis represents the net change difference relative to the baseline: biodiversity net change is the average weighted occurrence probabilities and carbon is Mg C. The outlined violin plot is identical across all scenarios, representing the distribution of values for the predicted net change between 2010 and 2020. Inside each intervention scenario’s violin plot, dots represent individual pixels that changed their forest type during this period. Dots are colored according to forest type: undisturbed primary forest (UPF), disturbed primary forest (DPF), secondary forest (SF), and deforested areas (D). In the predicted net change between 2010 and 2020, the dots naturally fit the shape of the violin plot, aligning with the net change distribution. However, in the other intervention scenarios, dots were constrained within the violin plot to highlight the net change difference between the predicted net change between 2010 and 2020 and the intervention scenarios. The maximum width of dot dispersal had to conform to the shape of the violin plot and high pixel densities around zero (i.e., areas of little to no net loss) resulted in a visual difference of positive contributions. The vertical dashed line represents the 2010 baseline, with values below the baseline indicating loss and values above representing gain.

Standardizing benefits by area allowed their effectiveness to be assessed independently of the spatial coverage (Fig. 3A-B). Results were broadly similar to unstandardized results for biodiversity: avoiding forest disturbances had the largest median direct benefit per unit area (0.0034 ± 0.0019 average weighted occurrence probability·ha^-1^·Yr^-1^), followed by avoided deforestation (0.0012 ± 0.0017), while restoration alone returned the lowest benefit per unit area (0.0006 ± 0.0012).

**Fig 3.**
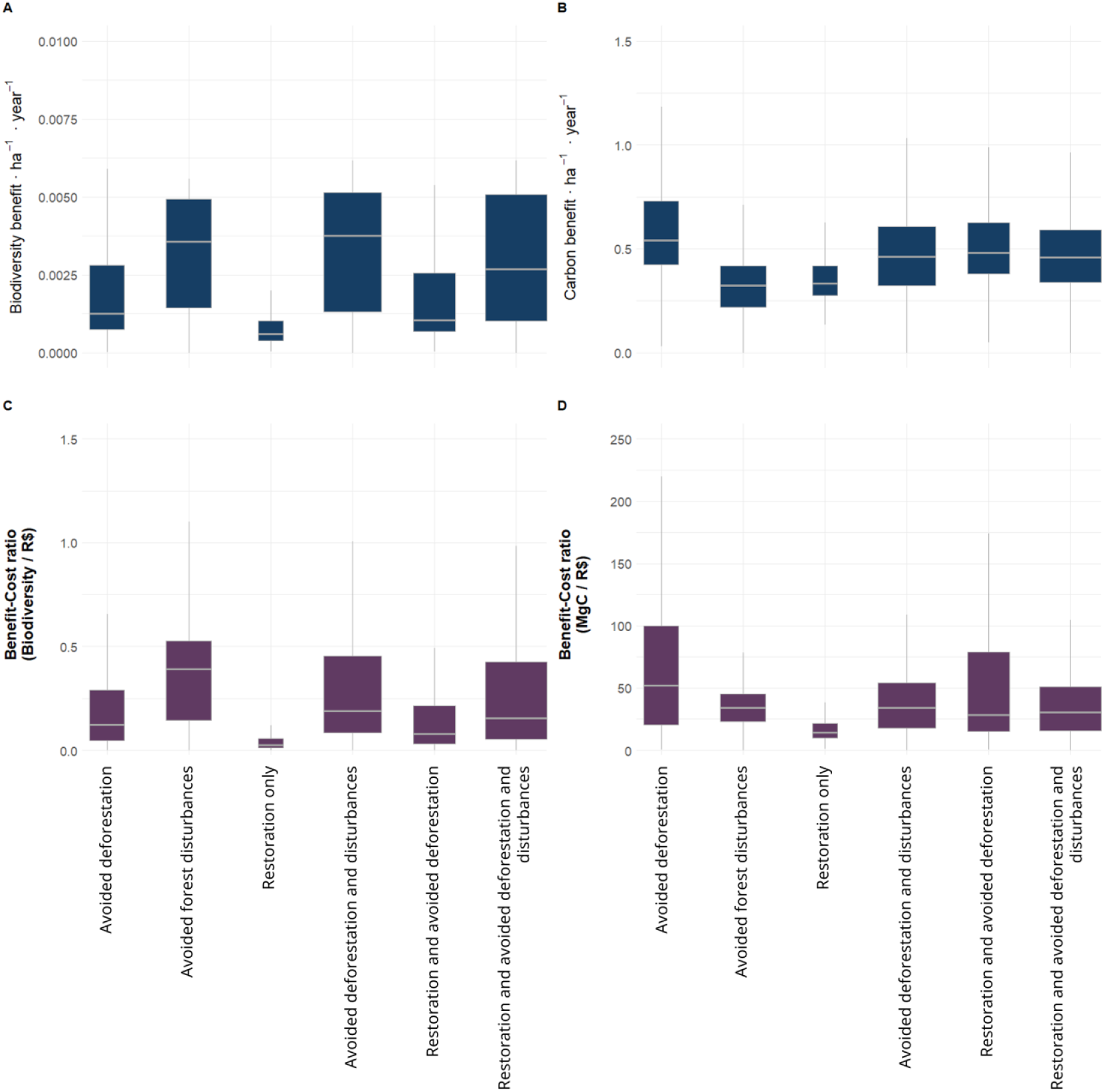
The per-area and per-cost benefits across the complete distributions of the intervention scenarios. Boxplots show: (A) and (B) benefits per hectare per year; (C) and (D) benefit–cost ratio (per R$10,000.00) for biodiversity and carbon, respectively.

For carbon, avoided deforestation provided the greatest median direct benefit (0.541 ± 0.233 Mg C·ha^-1^·Yr^-1^), while restoration and avoided forest disturbances returned lower outcomes (0.333 ± 0.103, and 0.319 ± 0.161, respectively). These direct effects are restricted to the pixels where interventions took place (i.e., the sites at which restoration occurred or where either deforestation or forest disturbance was prevented) and so do not estimate the additional benefits, such as protecting adjacent edges.

To guide policy implementation more broadly, we explored how the per-area biodiversity and carbon benefits covaried with property sizes and forest cover, both of which are central to environmental policy in Brazil (Table 1, Supplementary Material section 1.7, *26)*. The effectiveness of interventions was strongly associated with forest cover in 2010 (Fig. 4), with some of these patterns reflecting the land-use rules embedded in the intervention scenarios. For example, restoration becomes more effective than avoided deforestation at around 43% forest cover for biodiversity and 41% for carbon, close to the 50% forest legal reserve threshold. The lack of strong legal restoration requirements in small properties meant there were minimal benefits in these settings (fig. S16). Other property-level outcomes reflect the spatial distribution of deforestation and forest disturbance: avoided disturbances produced greater biodiversity gains than avoided deforestation in high-forest cover properties, reflecting that disturbances disproportionately affected intact, high-value forests, while deforestation tended to occur in more disturbed sites (fig. S15). Overall, these property level assessments demonstrate that the relative benefits of different interventions can be predicted at scale and modified by changes in property-level forest cover mandated by laws such as the NVPL.

**Fig 4.**
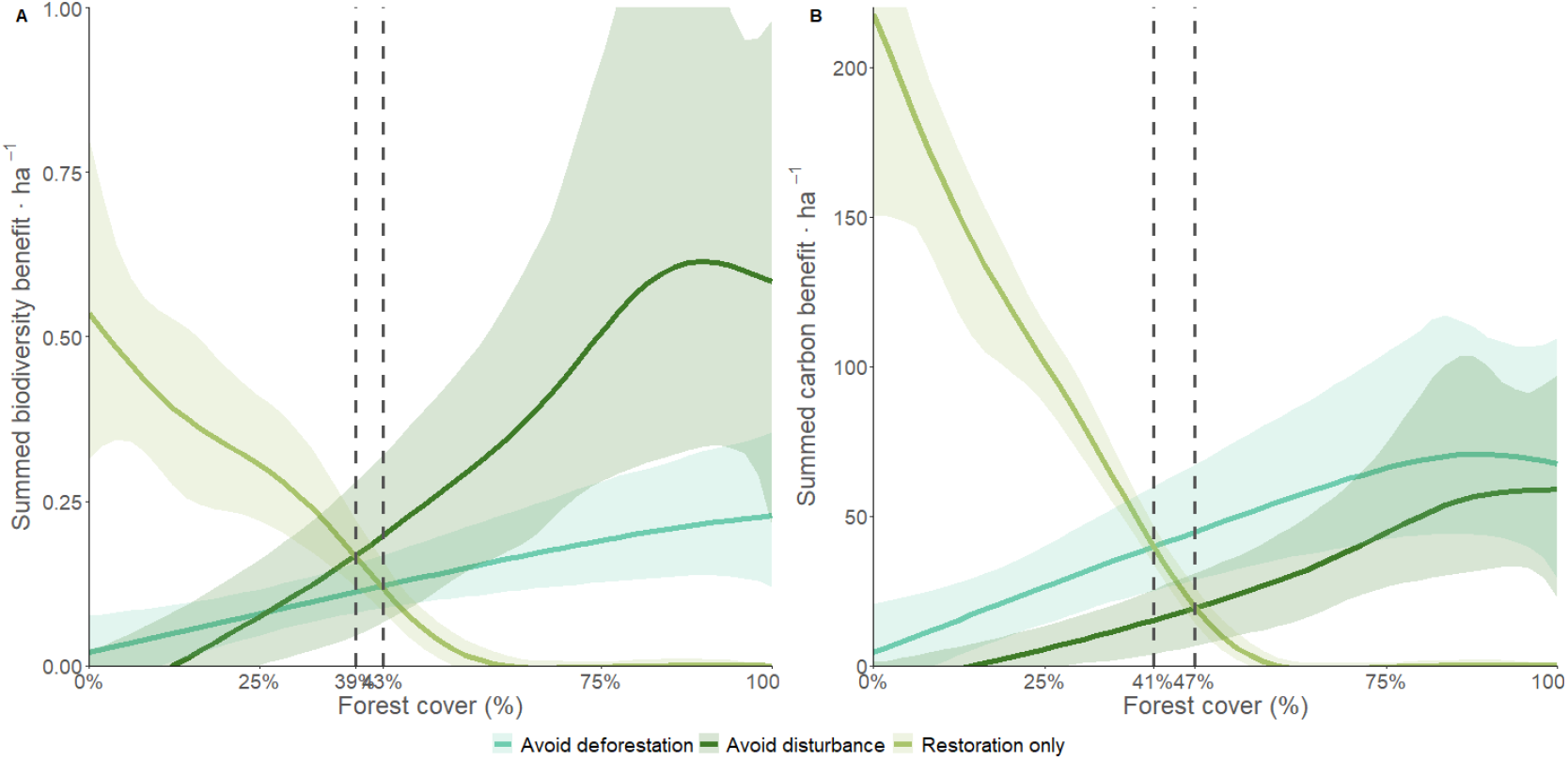
Variation in the summed value per-hectare of property of each intervention as a function of forest cover on medium-large properties. (A) Biodiversity benefits and (B) carbon benefits. Values were averaged across 1,000 bootstrap samples. Shaded areas show 95% confidence intervals. Vertical dashed lines indicate the crossover points where restoration begins to underperform avoided deforestation or avoided disturbances.

We next estimated the cost-effectiveness of each intervention, by dividing expected benefits by the estimated costs associated with implementation (fire control and restoration) and foregone opportunities (farm and logging incomes at the property level modelled from socio-economic interviews conducted across 499 properties and plot surveys [Supplementary Material section 1.8, figs. S17-S21, *26*]). Fire control was determined by the costs of establishing and maintaining firebreaks, whereas restoration (natural regeneration or active) was determined by the proximity to existing forest and land use type. Opportunity costs for farming were based on income lost due to agriculture and cattle raising operations, while for logging they based on income lost from wood harvesting (with limits set by Brazilian legislation). These were extrapolated spatially using the same RF modelling approach used for carbon stocks, adding two additional predictors: property size and distance to the urban center, representing socioeconomic, political and market variables (RMSE ±R$48.34 ha^-1^ year^-1^ and ±R$54.38 ha^-1^ year^-1^, for farming and logging income respectively). These were integrated into our interventions as follows: *avoiding deforestation* considered the foregone farming income; *avoiding forest disturbances* combined the costs of fire control and foregone logging income; *restoration* combined the costs of restoration and foregone farming income. Given the dynamic nature of economic conditions in Amazonia, we conducted a sensitivity analysis adjusting the key cost assumptions by –95% to +200%.

Introducing costs further reduced the value of restoration relative to avoidance intervention scenarios (Fig. 3C-D). For biodiversity, cost-effectiveness was greater for avoided forest disturbances (0.372 ± 0.234 average weighted occurrence probabilities per unit cost [R$10,000]), followed by avoided deforestation (0.123 ± 0.205), and restoration (0.025 ± 0.139). For carbon, avoiding deforestation returned the highest cost-effectiveness (52.0 ± 45.2 Mg C per unit cost), followed by avoided forest disturbances (33.8 ± 18.9) and restoration (14.2 ± 25.6). As it does not include foregone farming income, the predicted costs of avoiding forest disturbances (combined costs of fire control and foregone logging income) are relatively small and spatially less variable than other combinations (fig. S22). Avoided deforestation had the highest variance in benefit-cost ratio, reflecting high variation in opportunity costs among properties (fig. S22; *28)* as well as variation in benefits (Fig. 2). The low restoration cost-effectiveness reflects the high opportunity and implementation costs, although these were also highly variable depending on landscape context and restoration approaches. The relative cost-effectiveness of intervention scenarios was not sensitive to changes in the key cost variables (fig. S23). For example, the restoration remained less cost-effective than avoidance scenarios even if implementation costs were reduced by 95%, or if opportunity costs increased by over 100%.

Our counterfactual scenarios provide a heuristic exploration of the efficacy of three key interventions to conserve forest biodiversity and mitigate carbon loss, implemented individually and in plausible combinations. Headline results were broadly similar across benefit measures, geographic regions, and the exclusion of secondary forests in analysis (fig. S13, S14 and S22). It is important to note that our scenarios are not intended as literal representations of potential outcomes, but rather as diagnostic tools to understand the consequences of prioritizing different policy goals. Taken together, they provide four key policy-relevant insights.

## Avoiding forest disturbances is a highly cost-effective conservation strategy in consolidated frontiers

In both regions, avoided forest disturbances emerged as an essential and cost-effective strategy to tackle biodiversity loss and maintain carbon stocks in tropical forest frontiers, rivalling and, for biodiversity, even exceeding the benefits of avoiding deforestation. In part, this is due to their greater extent in our study regions, a pattern which is repeated across the entire Brazilian Amazon (6,*7,25)*. Our estimates of within-forest disturbances are also likely to be conservative, due to the challenges of detecting canopy disturbance (Supplementary Material section 1.3, 6,*26)*.

Avoiding forest disturbance therefore needs to become a central pillar of biodiversity and climate policies, requiring actions to prevent illegal or poorly managed selective logging, ensure authorized operations follow strict sustainability standards, and enable the adoption of effective firebreaks. These measures will benefit from regional initiatives that could not be costed in our spatially-explicit assessment, such as technical assistance to reduce ignition sources in agriculture and the funding of local fire brigades *(31)*. While the increasing dominance of fire-free mechanized agricultural practices could eventually reduce ignition sources *(32,33)*, this has not happened yet in our study regions: the extensive disturbances witnessed in the reference period were concurrent with the marked shift from pasture to mechanized soy *(34)*. However, pastures remain the predominant land use in the landscape, and there are important actors who rely on slash-and-burn practices *(28)*. The risks and intensity of forest fires will likely grow further due to climate change *(35,36)* and local disturbances such as logging and edge effects (6,*37)*. Importantly, our sensitivity analysis shows that avoided forest disturbances remains the most cost-effective intervention for biodiversity even under substantial increases in fire control costs (fig. S23).

### Avoiding deforestation remains a key priority in consolidated frontiers

Deforestation also continues in the consolidated frontier, with a loss of 95,193 ha of primary forest (5.3% of baseline primary forest) and 63,787 ha of secondary forests (42.6% of baseline secondary forest) between 2010 and 2020 across our study regions. The total impact of this continued forest loss was reduced by the prior impact of multiple disturbances (fig. S15), matching global patterns where disturbed regions show higher rates of deforestation (7,25). Avoiding deforestation would have much higher relative benefits at the earlier stages of frontier development where undisturbed forest is felled directly.

Nonetheless, deforestation remained the main driver of carbon loss, and an important driver of declines in biodiversity, and avoiding further deforestation in consolidated frontiers therefore remains a priority. This will require the enforcement of existing legislation, such as those mandating landowners to conserve APPs and RLs *(38)*, and incentives to avoid “legal” deforestation *(29)*. The latter can occur when properties have forest cover greater than the legal requirement, a situation that applies to *c*.32% of properties in our study regions, resulting in a “deforestable surplus” of *c*.157,300 ha. Public policies, such as the Brazilian National Payment for Ecosystem Services Law *(39)* or market mechanisms, such as those resulting from the Soy Moratorium (40) and the EUDR (18), could also play a crucial role in engaging farmers and associated commodity chains to comply with zero-deforestation commitments and broader environmental standards. Moreover, the greatest reductions in deforestation can be achieved in the 60 million ha of undesignated public lands that are vulnerable to land-grabbing and illegal activities in the Brazilian Amazon *(38,41)*. However, ongoing policy developments, such as Bill 2159/2021, which is currently being negotiated by lawmakers, could undermine these efforts by relaxing environmental licensing procedures, potentially opening new legal pathways for deforestation, even within protected areas *(42)*.

### Restoration cannot be a stand-alone strategy

Despite adding 80,705 ha of secondary forest to the regions, the biodiversity and carbon benefits of our restoration scenario were low (Fig. 1) and were swamped by the impacts of ongoing deforestation and forest disturbances (Fig. 2). Furthermore, implementing such large-scale restoration faces major challenges. From a legal perspective, our approach was overly optimistic as landowners could only achieve legal compliance by resolving the on-site deficits in their areas of APPs and RLs. In reality, most landowners prefer to compensate for these deficits by buying or leasing forested land elsewhere *(43)*. Moreover, there was large variation in the legal threshold requirements for deforestation, with this being much lower in Santarém than in Paragominas (figs. S13, S22), as well as in other municipalities in Pará state *(29)*. Going beyond legal compliance will therefore require incentive-based approaches, which must overcome high opportunity and implementation costs, the relatively low cost-effectiveness on a per area basis (Fig. 3), and the strong social “buy in” required for success *(44)*. Moreover, active restoration, which is often the focus of incentive-based restoration initiatives, poses other challenges when implemented at large spatial scales, such as limited seed availability and low seedling survival *(45)*.

Yet restoration can deliver the greatest benefits in areas with low levels of forest cover (Fig. 4, fig. S16) and provides synergistic benefits with avoidance strategies (Fig. 1.) showing it is an important part of a broader set of interventions. Our scenario also focused exclusively on restoring deforested areas, but the restoration of disturbed primary forests could offer a promising alternative *(46)*. Approaches such as biocultural restoration, which actively engage traditional communities and indigenous peoples, can also reduce costs and strengthen long-term socioecological resilience *(46)*. Finally, our ten-year assessment inevitably does not capture the full potential of secondary forests, which improve their carbon stocks and biodiversity similarity relative to primary forests over time *(47,48)*. However, it is doubtful that a longer-term assessment would have greatly improved the per-annum benefits or cost-effectiveness of secondary forests; the per ha benefits were annualized (Fig. 3), and the first ten years encompasses the period with the fastest rates of recovery for carbon and biodiversity *(47,49)*.

### Reversing biodiversity and carbon loss is only possible through combined interventions

Achieving net gains in biodiversity and carbon in frontier landscapes is more complex than many of the current market- and credit-based approaches which focus on single interventions such as restoration. We show that an integrated approach is required to reverse the losses of biodiversity and carbon and is the only way of unlocking important synergistic benefits (Fig. 1). These synergistic benefits – the net gains that exceed the additive effects of each intervention implemented individually – reflect a combination of indirect spatial effects and unmeasured interactions. The synergistic benefits derived from combining avoided deforestation and avoided forest disturbances reflect the high degradation risks in forests protected from deforestation: 32% of the deforestation in our study landscapes occurred in disturbed forests, and 57% within 500 m of a forest edge. However, in some cases, the benefits of avoiding forest disturbance can only be realized if forests are also protected from deforestation. Moreover, the benefits derived from combing restoration with avoided deforestation can be explained by buffering effects, with secondary forests adjacent to primary forests helping to reduce edge *(50)*. Such benefits were likely incentivized by selecting restoration locations based on proximity to primary forests, which, by encouraging natural regeneration, also reduces costs associated with seedling procurement, planting and maintenance, thereby boosting resource allocation for broader conservation efforts *(45,51)*.

Finally, although our analysis focused on biodiversity and carbon, the three key interventions are also likely to deliver regional benefits not assessed here. For instance, they can create economic opportunities and improve public health through cleaner water and reductions in air pollution and insect-borne diseases *(52-54)*. Forests also contribute to hydrological processes including long distance water transport, with benefits extending far beyond the Amazon and across South America’s grain belt *(55,56)*. These additional benefits are important, but further research is needed to understand the relative contribution of intact primary, disturbed primary and secondary forests to these flows *(57)*.

The first ever UNFCCC COP in the Amazon places renewed focus on the measures required to safeguard Amazonian ecosystems and the well-being of its people. Our results show that these discussions need to place much greater emphasis on actions to avoid forest disturbances, which is both cost-effective and delivers the greatest summed benefits for biodiversity. Furthermore, we caution that the growing interest and focus on restoration risks having only a minimal impact unless integrated with measures that avoid further deforestation and forest disturbances. Crucially, we show that it is possible to halt and reverse biodiversity and carbon losses across the vast tropical forest frontier landscapes, but that combined interventions are required to unlock the greatest gains.

## Acknowledgments

We are grateful to our numerous field and laboratory assistants, particularly to our parabotanists Nelson Rosa *(in memorian)* and Manoel Cordeiro. We also thank the farmers and workers unions of Santarém, Mojuí dos Campos, Belterra and Paragominas, and all collaborating private landowners for their support. This is paper #128 of the Rede Amazônia Sustentável publication series.

## Author contributions

Conceptualization: LSM, JRT, JB, JF, ACL, RM and TAG; Methodology: LSM, JRT, JB, GDL, JF, ACL, EB, RM, LEOCA, PHSB, SFBF, RDG, PGM, NGM, SSN, LP, JS, ICGV, CV and TAG; Investigation / Analyses: LSM, JRT; Visualization: LSM, JRT; Writing – original draft: LSM, JRT, JB; Writing – review & editing: All.

## Data and materials availability

To ensure the full reproducibility and transparency of our research, all the post-processed data and scripts/codes developed in this study are made publicly available and briefly described to facilitate reproducibility and applicability on a GitHub repository (https://github.com/miralaba/conservation_opportunities).

## Supplementary Materials

Materials and Methods

Figs. S1 to S23

Tables S1 to S2

References *(57–90)*

