## Supplementary Material for "Combined interventions are required to reverse biodiversity and carbon losses in tropical forest frontiers"

Science
AAAS

Supplementary Materials for

### Combined interventions are required to reverse biodiversity and carbon losses in tropical forest frontiers

Leonardo S. Miranda, James R. Thomson, Joice Ferreira, Toby A. Gardner, Erika Berenguer, Alexander C. Lees, Ralph Mac Nally, Luiz E. O. C. Aragão, Pedro H. S. Brancalion, Silvio F. B. Ferraz, Rachael D. Garrett, Paulo G. Molin, Nárgila G. Moura, Sâmia S. Nunes, Luke Parry, Juliana M. Silveira, Ima C. G. Vieira, Cecilia Viana, Jos Barlow

**The PDF file includes:**

Materials and Methods

Figs. S1 to S23

Tables S1 to S2

References (*58-90*)

Contents


**1. Materials and Methods** 4

***1.1. Study area*** 4

*1.2.* ***Remote sensing datasets*** 5

*1.3.* ***LULC change and classes*** 6

***1.4. Interventions and Scenarios*** 7

***1.5. Restoration candidate areas*** 8

*1.5.1. Brazil's Native Vegetation Protection Law and the legal requirements for restoration* 8

*1.5.2. Estimating legal forest deficits in the study regions* 9

*1.5.3. Identifying areas for restoration* 10

*1.6.* ***Biodiversity, carbon and socio-economic surveys*** 11

***1.7. Spatially explicit biodiversity and carbon benefit metrics*** 11

*1.8.* ***Spatially explicit costs*** 13

**2. Supplementary Figures** 16

***Figure S1*** 16

***Figure S2*** 17

***Figure S3*** 19

***Figure S4*** 20

***Figure S5*** 21

***Figure S6*** 23

***Figure S7*** 25

***Figure S8*** 26

***Figure S9*** 27

***Figure S10*** 28

***Figure S11*** 29

***Figure S12*** 30

***Figure S13*** 31

***Figure S14*** 33

***Figure S15*** 35

***Figure S16*** 36

***Figure S17*** 37

***Figure S18*** 38

***Figure S19*** 39

***Figure S20*** 40

***Figure S21*** 41
***Figure S22*** 42
**Figure S23** 44
**3. Supplementary Tables** 45
Table S1 45
Table S2 48
**4. References and Notes** 70

#### 1. Materials and Methods

##### *1.1. Study area*

Our study was conducted in Pará, a Brazilian state in the eastern Amazon (fig. S1), known for its exceptional biodiversity but also a major source of deforestation and degradation (*8*). We focused on two regions of Pará, including four municipalities: Paragominas (PGM); and Santarém (only the portion of the municipality on the right bank of Tapajós River), Mojuí dos Campos and Belterra. The later three share boundaries, so we will refer to this region as Santarém (STM) from now on. This cut for the STM region has the purpose of maintain consistency with the main biogeographic endemism areas, delimited by the largest tributaries of the Amazon basin, and to match the study area target of several studies conducted by the Sustainable Amazon Network (see also *Biodiversity, carbon and socio-economic surveys* section below). Paragominas and Santarém are located in distinct biogeographic endemism areas, Belém and Tapajós (*sensu* Silva et al., *58*) respectively, have differing histories of human colonization, and different experiences with environmental governance (*28*).

Paragominas is located in the easternmost endemism area of the biome, which is recognized as a biodiversity hotspot with high levels of species richness and endemism, particularly in birds and plants. The area harbors several taxa that are critically endangered due to habitat loss, making them particularly vulnerable to deforestation and fragmentation (*59,60*). Despite PGM still maintaining more than two-thirds of its native forest cover (fig. S2), the neighboring area has already lost over 70% of its original forest cover, primarily due to agricultural expansion, logging, and cattle ranching (*61*). The municipality was founded in 1959, which means it has a relatively recent history of land use and human occupation compared to many Amazonian municipalities. It experienced a significant influx of settlers during the 1950s and 1960s, driven by cattle ranchers migrating from southern Brazilian states, and during the 1980s and 1990s with the boom of the timber industry. In the early 2000s, large-scale mechanized agriculture became a prominent land use (*28*). This relatively recent and fast history of human occupation and land conversion placed PGM on the government's "deny list" of municipalities due to the extremely high deforestation rates, which restricted access to financial credit until deforestation levels were reduced. However, the region took steps towards sustainability, particularly through the *Município Verde* (Green County) initiative, which aimed to promote sustainable land-use systems (*62*). As a result of this background, PGM is more comparable to other fragmented regions in the southern Amazonia's Arc of Deforestation, where historical deforestation and land conversion have caused an increase in edge effects and decreased connectivity, leaving remaining forests vulnerable to degradation threats like wildfires. This has also resulted in the loss of ecological value of small, isolated patches, escalating the vulnerability to biodiversity.

Biogeographically, STM is positioned within an endemism area that encompasses a vast portion of the central Amazon and remains more intact than PGM (fig. S2), though degradation is increasing due to prevalent wildfires, illegal logging and mining, and infrastructure development (*29*). This area supports a distinct set of endemic species, and the biodiversity here is influenced by the region's more humid climate and extensive river systems (*58*). Many species in this area are highly specialized and depend on continuous, old growth primary forest, making them particularly vulnerable to degradation (*58,60*). Santarém is one of the oldest cities in the Amazon, founded in 1661, has a long history of human settlement and, together with the neighboring municipalities of Belterra and Mojuí dos Campos, have been densely populated by small farmers for more than a century (*28*). The most recent land use and human occupation in this region has been closely linked to the construction of federal highways, which facilitated human colonization and agricultural expansion, with the emergence of large-scale mechanized agriculture also in the early 2000s. Therefore, the STM shares characteristics with the more intact central regions of the Amazon where threats of degradation and deforestation are increasing and becoming more widespread.

These regions in Pará exemplify the interplay between biodiversity, resource exploitation, and sustainability practices, reflecting broader Amazonian dynamics and offering insights into the complexities of conservation amid environmental change.

##### *1.2.* ***Remote sensing datasets***

The remote sensing datasets used as explanatory variables in this study are derived from sources that are openly available. Collection 8 of the Mapbiomas project was utilized for capturing Land Use – Land Cover (LULC) from 2007, 2010 and 2020 (available at https://brasil.mapbiomas.org/en/colecoes-mapbiomas/). The age of secondary forest in 2010 and 2020 was processed using Silva Jr et al. (*63*) (available at https://github.com/celsohlsj/gee_brazil_sv). Time since last disturbance (fire scars and logging) was processed using Mapbiomas fire collection 2 from 1985 to 2020 (available at https://brasil.mapbiomas.org/en/colecoes-mapbiomas/), and vectorized data from the monitoring system of deforestation and degradation of the Brazilian National Institute for Space Research (INPE, acronym in Portuguese): DEGRAD from 2007 to 2016 (available at http://www.obt.inpe.br/OBT/assuntos/programas/amazonia/degrad/acesso-ao-dados-do-degrad) and DETER-B from 2016 to 2020 (available at http://terrabrasilis.dpi.inpe.br/file-delivery/download/deter-amz/shape). Mean annual temperature and precipitation from to 2010 and 2020 were derived from the monthly land surface temperature and rainfall archived on NASA Earth Observations (available at https://neo.gsfc.nasa.gov/). The elevation, river and roads were taken, respectively, from SRTM (available at https://earthexplorer.usgs.gov/), HydroSHEDS (available at https://www.hydrosheds.org/), and the Imazon institute. Finally, data on rural properties were downloaded from Brazilian National System of Rural Environmental Registration (SISCAR, acronym in Portuguese; available at https://www.car.gov.br/). All environmental layers were downloaded in the highest spatial resolution that was available – 30m in most cases. In cases where the resolution was less than 30m, a resample procedure was performed.

##### *1.3.* ***LULC change and classes***

The steps for processing input data, redefining LULC classes for 2010 (baseline) and 2020 (observed changes), and building landscapes relative to each intervention scenario are summarized in fig. S3 and detailed below. We first distinguish between forest (the "forest formation" class) and deforested areas (including "pasture", "soybean" and "other temporary crops" classes) using the LULC maps from Mapbiomas. Those areas under permanent forest cover and without signs of human modification (deforestation, fire and/or logging) were defined as Undisturbed Primary Forests (UPFs). Secondary forests (SFs) were defined as forests recovering after complete vegetation removal, where the age of secondary forests was obtained from time-series analysis of LULC maps from Mapbiomas according to Silva Jr et al. (*63*). Here we also distinguish between Undisturbed Secondary Forest (uSF) and Disturbed Secondary Forest (DSF, those with fire scars). We created a time series using maps from Mapbiomas fire, and shapefiles from DEGRAD and DETER-B of the Brazilian degradation monitoring system (INPE), to calculate the time since last disturbance event. Thus, areas under permanent forest cover but presenting fire scars and/or selective logging were defined as Disturbed Primary Forests (DPFs), with the distinction between those that have been disturbed only once (uDPF) and those that have been disturbed multiple times (RDPF). To estimate the time since the last disturbance as accurately as possible, we combine the three datasets yearly separately. Although the three datasets do not match the entire time-series, they are still very effective in capturing fire scars. However, only the DEGRAD and DETER-B systems detect signals of selective logging. It is important to mention that there are some limitations in relation to DEGRAD that were based on lower resolution LANDSAT and CBERS images compared to the current DETER-B monitoring system, which is based on WFI and CBERS-4 sensors (*64*). Therefore, we acknowledge that selective logging is probably underestimated.

##### *1.4. Interventions and Scenarios*

To assess the effects of different conservation interventions on biodiversity and the carbon stocks, and their economic costs, we simulated several scenarios based on the observed trajectories of deforestation, degradation and restoration between 2010 and 2020; as well as policies, laws and programs that are being implemented in the Brazilian Amazon (Table 1). For instance, policies have mostly focused on reducing deforestation and promoting forest restoration, notably the Action Plan for the Prevention and Control of Deforestation in the Brazilian Amazon (PPCDAm, *14*) being particularly effective in its earlier phases (2007– 2010). In addition, private rural areas are regulated by Brazil's Native Vegetation Protection Law (NVPL, *13*), which mandates the retention of up to 80% forest cover in the Amazon as Legal Reserves and Permanent Protection Areas, as well as the restoration of illegal deforestation to reach the thresholds defined by law (see more details below). Restoration is also incentivized by Brazilian agricultural credit schemes as part of their Low Carbon Agriculture Plan (*65*), REDD-related "Floresta+" funds, and national and international commitments, such as the National Plan for Native Vegetation Recovery (Planaveg; 12 million hectares goal; *20*), the Bonn Challenge (*19*), the Paris Agreement (*66*) and the Global Biodiversity Framework's Target 2 (although awaiting Brazil's National Biodiversity Strategies and Action Plan). Moreover, land abandonment following deforestation has led to the forest regrowth of over 18 million ha in the Brazilian Amazon (*24,63,67*), with much of this on private properties or undesignated lands (*68*). However, forest degradation from fires, selective logging, and other disturbances remains an increasingly significant concern and continues to be marginally addressed (*6,21-25,69*).

Therefore, we recreated landscapes that simulate the following scenarios: (i) Avoiding deforestation; (ii) Avoiding within-forest disturbances, (iii) Restoration only, (iv) Avoiding both

– deforestation and disturbances; (v) Restoration and avoid deforestation; and (vi) Restoration and avoid both (figs. S5-S6). The LULC layers were recreated over a 10-year window where, for instance, deforestation from 2010 to 2020 was avoided (scenario i); or disturbances from fire and logging was avoided in that period (scenario ii); or the deficit of legal reserves (the legal compliance with the NVPL) were restored, resulting in secondary forests with 10 years (scenario iii; see next section); or combinations of these main strategies (scenarios iv-vi). We also consider that these conservation interventions were applied to either primary and secondary forests, or to primary forests only. Although we acknowledge that these are not the only possible scenarios or even the most realistic (zero deforestation, or zero degradation, or 100% legal compliance with restoration), our goal is to compare them as a heuristic exercise to inform policy and to establish benchmark outcomes against which future or more realistic policy pathways can be assessed.

##### *1.5. Restoration candidate areas*

###### *1.5.1. Brazil's Native Vegetation Protection Law and the legal requirements for restoration*

The Brazil's NVPL (*13*) establishes a complex framework for managing private lands, balancing environmental protection with economic activities. Rural properties are, therefore, divided into two main areas, one for production and the other for conservation and sustainable management of resources. This conservation area is further categorized into Permanent Protection Areas (APPs, acronym in Portuguese) and Legal Reserves (RL, acronym in Portuguese). The first are designed to protect sensitive habitats like riparian zones, steep slopes and hilltops, permitting only low-impact activities such as ecotourism. The former are meant for biodiversity conservation and/or sustainable use, allowing controlled economic activities such as selective logging, but prohibiting deforestation. The size and extent of APPs are determined by the width of adjacent watercourse. For example, small streams (up to 10 m wide) require a riparian buffer zone of 30 m on each side. As the width of the watercourse increases, so does the required width of the riparian buffer, which can range up to 500 m on each side for very large rivers (>500m). It is worth noting that APPs are not separate but are part of the RL of a property. The size of RL, in turn, is determined by several factors, including its location, property size and when deforestation occurred. For instance, if the property is located in the Amazon, the RL area can be up to 80% of its size; or if the property is less than four fiscal modules (a measure used by the Brazilian government to classify a rural property, varying from 5 to 110 hectares depending on the municipality), the RL area may be equivalent to the existing native vegetation cover on July

2008. On the other hand, specific conditions allow for the reduction of RL areas, such as when more than 50% of a municipality is in Protected Areas (Conservation Units and Indigenous Lands), or for properties located in areas indicated by an Ecological and Economic Zoning plan.

Importantly, the NVPL sets specific rules for the restoration of RL, with requirements varying based on watercourse width (for APPs), property size and the timing of deforestation. For instance, properties that have cleared vegetation in APP after July 2008 are obligated to fully restore these areas to the width specified by the NVPL. The property size also influences the restoration requirements. For properties smaller than four fiscal modules, the RL is set at the level of native vegetation cover in July 2008. Small properties that have cleared vegetation after July 2008 are required to restore to the same amount of vegetation on that date. Medium-large properties (more than four fiscal modules) that have cleared more than 50% of their forest are required to restore or compensate up to 80% of it. Therefore, the NVPL's detailed criteria for restoration were designed to cater to different scenarios, supposedly to ensure a balanced approach between environmental conservation and land-use.

###### *1.5.2. Estimating legal forest deficits in the study regions*

Our initial data included 10,370 properties, 42 agrarian reform settlements and six protected areas, which resulted in 10,418 instances registered in the study area (2,645 in PGM and 7,773 in STM). It is important to note that numerous errors and disputes over properties boundaries, largely because the self-declared nature of provisional CARs, allow inconsistencies in property extents and, consequently, overlaps. To address these discrepancies and avoid double counting, we adopted a series of exclusion criteria, following Nunes et al. (*29*): (i) in cases where properties were duplicated within the system, evidenced by identical polygons and sizes, one of the duplicates was randomly excluded; (ii) for cases where the overlap between two properties exceeded 80%, the smaller of the two properties was excluded; (iii) properties overlapping more than 30% with agrarian reform settlements were excluded from the analysis; and (iv) we also excluded properties that had more than 50% of their area overlapping with water bodies or areas not suitable for registry, such as protected areas. In addition, properties with less than 0.1 ha were also excluded and our restoration scenario did not include protected areas. After the exclusions, our data contained 6,900 properties (1,539 in PGM and 5,361 in STM). Property sizes were recalculated from the SISCAR polygons, and these were expressed in terms of fiscal modules, which vary by municipality. In our study, one fiscal module is equivalent to 55 ha in

PGM and 75 ha in STM, and this difference is fundamental in the classification of properties by the NVPL and in the determination of restoration requirements. The existing forest cover inside the properties was estimated as a percentage of the total property area by using the "forest formation" class of the LULC maps for the years 2007 and 2010, to identify the properties with legal deficits of forests. The legal requirements for restoration are contingent on the width of the watercourse, property size, and when deforestation occurred, as per NVPL. The width of the watercourse in meters was estimated using HydroSHEDS data, which contained the attributes of watercourse's length and area. Hence, based on the estimated width of watercourse, we delimited the areas of APPs by property, verified those without forest cover between 2007 and 2010, and defined them as having forest deficit. For medium-large properties (≥4 fiscal modules), those with less than 50% forest cover in 2010 were defined as having forest deficit. Whereas, for small properties (<4 fiscal modules) or those classified as agrarian reform settlements, the determination of the forest deficit was based on the comparison of forest cover between 2007 and 2010. If there was a decrease in forest cover or if the forest cover was below 50%, these properties were also defined as having forest deficit.

*1.5.3. Identifying areas for restoration*

We developed a framework to hierarchically allocate areas that could be restored under the current legal requirements by integrating spatial data on property boundaries, forest deficits and distances to existing forest (fig. S4). First, since restoration of APPs is mandatory, all forest deficit associated with APPs were designated as candidate areas for restoration. Then, considering that APPs comprise the RL, we assessed the properties where the RL remained below the legal limits after potential APP restoration, and how much forest each one required to restore to meet RL compliance. Finally, we determined the spatial allocation of the remaining forest deficit within the properties based on the distance to the existing forest. Unlike APP restoration, there are no legal regulations specifying where RL should be allocated, so we used a strategy based on ecological principles (*50*) as well as evidence-based costs and success (*45*). While we recognize this will not be the only criteria used when restoration decisions come to be made at the property level, it at least gives a first order approximation of where to promote natural regeneration, enhance ecological connectivity and conserve biodiversity.

All procedures described so far were developed in the R environment (v4.3.0; *70*), based mainly on spatial data processing functions using raster (v3.6.20; *71*), sp (v1.6.1; *72*) and sf (v1.0.13;

*73*) packages as well as custom functions created using base package.

##### *1.6. **Biodiversity, carbon and socio-economic surveys***

Standardized field surveys were undertaken in both PGM and STM between 2010 and 2011. The study regions were divided into drainage catchments, from which thirty-eight were selected (mean size ~5000 ha), representing the gradient of forest cover in 2010 (the baseline). Using a stratified random sampling design, between eight and twelve transects of 300×10 m was allocated to each catchment (208 in PGM and 173 in STM), on "*terra firme*" habitat, and separated by >1.5 km to minimize spatial autocorrelation. Two repeated 15-min bird surveys were undertaken at three sampling points within these transects (0, 150, 300 m). We performed point counts between 15 mins before dawn and 0930. We also measured and identified at the species level all trees and palms ≥10 cm diameter at breast height (DBH) in plots of 10×250 m, and smaller individuals (2 ≤ DBH <10) in five subplots of 5×20 m. Aboveground carbon values were assumed to be 50% of total biomass (*74*), which was calculated from individual DBH, species-specific wood density (*75*), and allometric equations (*76*).

Socioeconomic surveys were conducted in 499 properties (171 in PGM and 328 in STM), which were spatially congruent with the terrestrial biodiversity data (within the same catchments). Data on production and price values were reported, encompassing annualized sales of agricultural products and cattle ranching as well as operating costs of property maintenance, machinery, input (e.g., seeds, fertilizer, feed) and wage labor. For further details regarding the survey design, see Gardner et al. (*27*).

##### ***1.7. Spatially explicit biodiversity and carbon benefit metrics***

Biodiversity and aboveground carbon values were predicted using Random Forest (RF) models trained and tested on data collected in the surveys described above. These models allowed prediction of occurrence probabilities for individual species and carbon stocks conditional on forest type (undisturbed primary forest, disturbed primary forest, secondary forest), forest condition (forest age, time since last disturbance, edge densities) and landscape context (distances from rivers and roads, climate, elevation). These environmental predictors were derived from the related environmental data layers also described above, and referring to the 2010 baseline (table S1). A total of 24 predictor variables were obtained, which were tested for collinearity using Pearson's correlation tests and the variation inflation factor through the functions *removeCollinearity()* of the virtualspecies package (v1.5.1; *77*) and *vifcor()* of the usdm package (v1.1.18; *78*). After this assessment, the models were fitted using 14 variables (fig. S7). Occurrence records of birds and plant species and the estimated carbon stock in the plot were used as response variables. In RF modelling, there are three key parameters: the number of trees (*ntrees*), the number of variables at each tree node (*mtry*) and the minimum number of terminal nodes (*nodesize*), which were optimized using a grid search to explore an array of *ntree* and *mtry* values. Specifically, the number of *ntrees* varied between 100 and 2000, with an increment of 50; and the number of *mtry*, between 3 and 14, with an increment of 1. Default values for *nodesize* were used. To ensure robustness in our model predictions, we adopted a "*k*-fold cross validation", with individual catchments iteratively held out and used to test predictions from models trained on remaining catchments. The final models were evaluated using the area under receiver operator curves (AUC), for biodiversity; and root mean square error (RMSE), for carbon. In addition, the importance of each variable within the final model was quantitatively assessed, by calculating the mean decrease in accuracy (%IncMSE). This indicator measures the percentage increase in the mean square root when a specific variable is omitted from the predictors set. The final models were used to project biodiversity and carbon outcomes for each scenario. We implemented this analysis in R using the randomForest (v4.7.1.1; *79*) and caret (v6.0.94; *80*) packages.

For biodiversity data, we focus our modelling approach on forest-dependent species. Our list included species with at least 40% of their occurrence records in survey sites with more than 80% of undisturbed primary forests within a 1km buffer (initially 242 birds and 909 plants; table S2). We also filtered each species' records to obtain the maximum number of occurrences that were at least 1.5 km apart, which is the minimum distance between transects, using the spThin package (v0.2.0; *81*). This procedure reduces sampling bias and improves model performance (*82*). However, after this filtering process, we were unable to apply our modelling approach for several species due to a low number of records (<10 records). Thus, our final biodiversity database consisted of 160 bird species and 426 plant species. The reduction in the number of species analyzed did not change the species' representativeness in terms of the range size (fig. S8). Additionally, to account for variation in species conservation value, we weighed the occurrence probability by the species' geographic range size, which is a good indicator of species threat status and is correlated with extinction risk (*83*). Range sizes were taken from the

BirdLife Data Zone (*84*) and the World Checklist of Vascular Plants (*85*). We applied a weighing, in which we gave the species with the smallest range size a conservation value of 0.999; the one with largest range size a value of 0.001; and all other values were mapped linearly between these extremes (*8,48*).

Once these spatially explicit values were predicted for each pixel, we used them to calculate net changes in biodiversity and carbon between 2010 and 2020 under both the observed land-use trajectory and each counterfactual intervention scenario. Net change was defined as the sum of all pixel-level differences in predicted values across the landscape, comparing the state at the baseline (2010) and end (2020) of the period. We then derived benefits for each intervention scenario by comparing the predicted net change under that scenario to the "observed" net change. Specifically, the benefit of a scenario was calculated as:

*Benefit = Net change under observed 2020 – Net change under intervention scenario*

All benefit metrics were computed per ha and aggregated by region and intervention type, allowing for both spatially explicit comparisons and cost-effectiveness assessments.

To assess whether the relative benefits of interventions are consistent across varying contexts beyond our study regions, we implemented a bootstrap analysis using properties as the unit of analysis. Using 6,900 properties, we calculated the summed carbon and biodiversity benefits per ha of property and classified each property by size class (small and medium-large) and forest cover in 2010 (0–1 scale), using these as proxies for different frontier conditions. We performed 1,000 iterations (100 subsamples of 20% of the dataset, with 100 replicates each) to estimate the mean benefit as a function of the property characteristics under each intervention scenario. Confidence intervals (95%) were derived from the bootstrapped distributions. This approach provides a measure of how robust intervention outcomes are to spatial heterogeneity within Amazonian frontier areas.

##### *1.8.* ***Spatially explicit costs***

The costs associated with each scenario were estimated using implementation costs (fire control and restoration) and opportunity costs (foregone logging income and foregone farm income). Forest fire control costs were estimated as the cost of creating and maintaining fire breaks at forest edges within properties. According to Institute for Environmental Research of the Amazon (IPAM, acronym in Portuguese; *86*), it would take on average three days to clear 100 meters of fire break (~33.333m/day) costing R$100 per day; and re-cleaning would need to occur every four years on average. Therefore, the estimated forest fire control costs per hectare per year were *((L / 33.333) x 100) / PS / 4*; where *L* is the length in meters of forested perimeter within a property and *PS* is the property size.

The costs of restoration are largely determined by past land use and distance from remaining natural habitats, so we estimate the costs associated with three strategies according to Brancalion et al. (*45*): (i) natural regeneration without a fence, R$189.13 /ha; (ii) natural regeneration with fence, R$1,331.55 /ha; and (iii) active restoration, R$7,899.71 /ha. Natural regeneration was attributed to pixels 500 m or less from the forest. If they were pastures, we attributed natural regeneration with fence, otherwise without a fence. We attributed active restoration to pixels that are more than 500 m away from the forest.

Regarding the foregone logging income, we first estimated the species-specific timber values as the product of the minimum market price of one $m^3$ (Interno\$/$m^3$; established by the Pará state Finance Secretariat, *87*) and tree volume. The later was calculated with the allometric equation: *Volume=exp(-6.86 + 1.994log(DBH))*; according to Vidal et al. (*88*). We then estimated a maximum potential value for logging revenues on surveyed transects as the sum of the most valuable trees, ≥40 DBH, and up to a maximum harvestable volume of 30 $m^3$/ha, considering current Brazilian forest management regulations, which allow a logging intensity of 0.86 $m^3$/ha/year in a 35-year cutting cycle (*89*). The overall planning and operating expenditures of logging activities were estimated at R\$66.23/$m^3$ by Imazon (*90*) in the PGM region, so we consider the net logging income as the difference between the maximum potential revenue of logging and the expenditures.

Similarly, net farm income was calculated by subtracting farm expenditures from farm revenues. Finally, we modelled income per hectare per year of logging and farming, using the same environmental predictors as for biodiversity and carbon, but adding two variables: property size and distance to the urban center (table S1). We also followed the RF modelling approach that was used for carbon.

The total final costs for each scenario were taken as: (i) the foregone farm income for avoiding deforestation; (ii) fire control AND foregone logging income for avoiding within-forest disturbances; (iii) restoration AND foregone farm income for restoration only; (iv) fire control

AND foregone logging AND farm income for avoiding both; (v) restoration AND foregone farm income for restoration and avoid deforestation; and (vi) restoration AND fire control AND foregone logging AND farm income for restoration and avoid both.

We acknowledge that conservation interventions also involve other costs. However, there is a lack of quantitative data on other types of economic costs, or even political and social costs, to incorporate into our modelling approach. The modelling framework we used for the opportunity costs had low explained variance and high uncertainties (see figs. S16 and S18), which is likely due to the non-market/environment-related predictor variables used. Although we included critical drivers such as property size and distance to the urban center, both identified as the most important variables in our analysis, other market variables such as the international fluctuation of commodity prices could potentially better explain the opportunity costs of logging and farming. Nevertheless, the complexity and variability of market dynamics and socio-political context in the Amazon make it difficult to identify and measure the full range of influencing factors.

Recognizing the uncertainty in implementation and opportunity costs, we tested whether our benefit-cost ratios findings were sensitive to key cost assumptions. Each cost component was independently varied from 95% cheaper to 200% more expensive in increments of 10%. For every cost adjustment, we applied 100 bootstrap iterations to each of 100 randomly selected subsamples (N = 200,000 pixels per subsample), recalculating the mean benefit per cost unit (i.e., R$10,000) for carbon and biodiversity. This design allowed us to identify thresholds at which the rank order of cost-effectiveness could change, thus assessing how resilient our conclusions are to plausible economic variability.

#### 2. Supplementary Figures

***Figure S1***

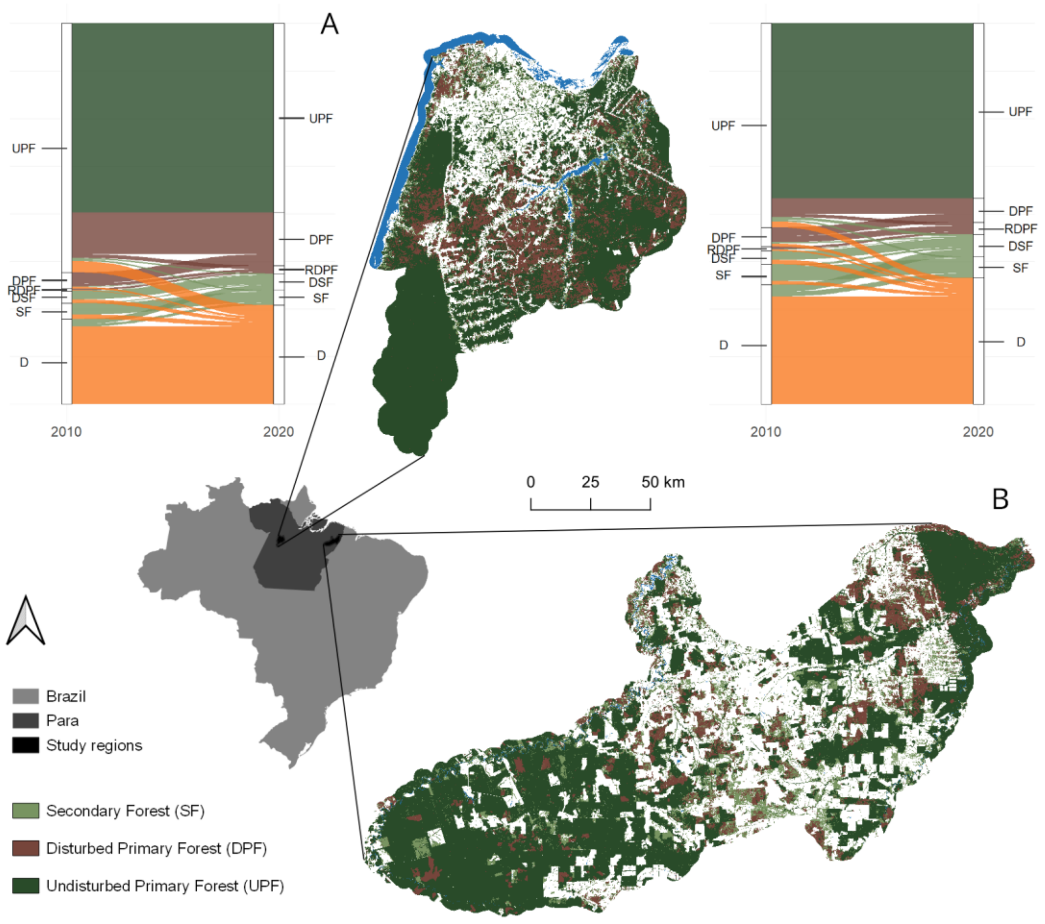


Fig. S1.

**Forest cover transitions and spatial distribution of forest types in the two study regions in Pará, Brazil.** Map of Brazil showing the state of Pará and the location of the two study regions: Santarém (center-top) and Paragominas (bottom-right). Forest types in 2020 are shown in dark green: Undisturbed Primary Forest (UPF), brown: Disturbed Primary Forest (DPF), light green: Secondary Forest (SF). Sankey diagrams illustrate the forest cover transitions between 2010 and 2020 in Santarém (top left) and Paragominas (top right). Despite identical color codes, different disturbance trajectories are distinguished in the Sankey diagrams: Recurrently Disturbed Primary Forest (RDPF) and Disturbed Secondary Forest (DSF) and Deforestation (D).

*Figure S2*

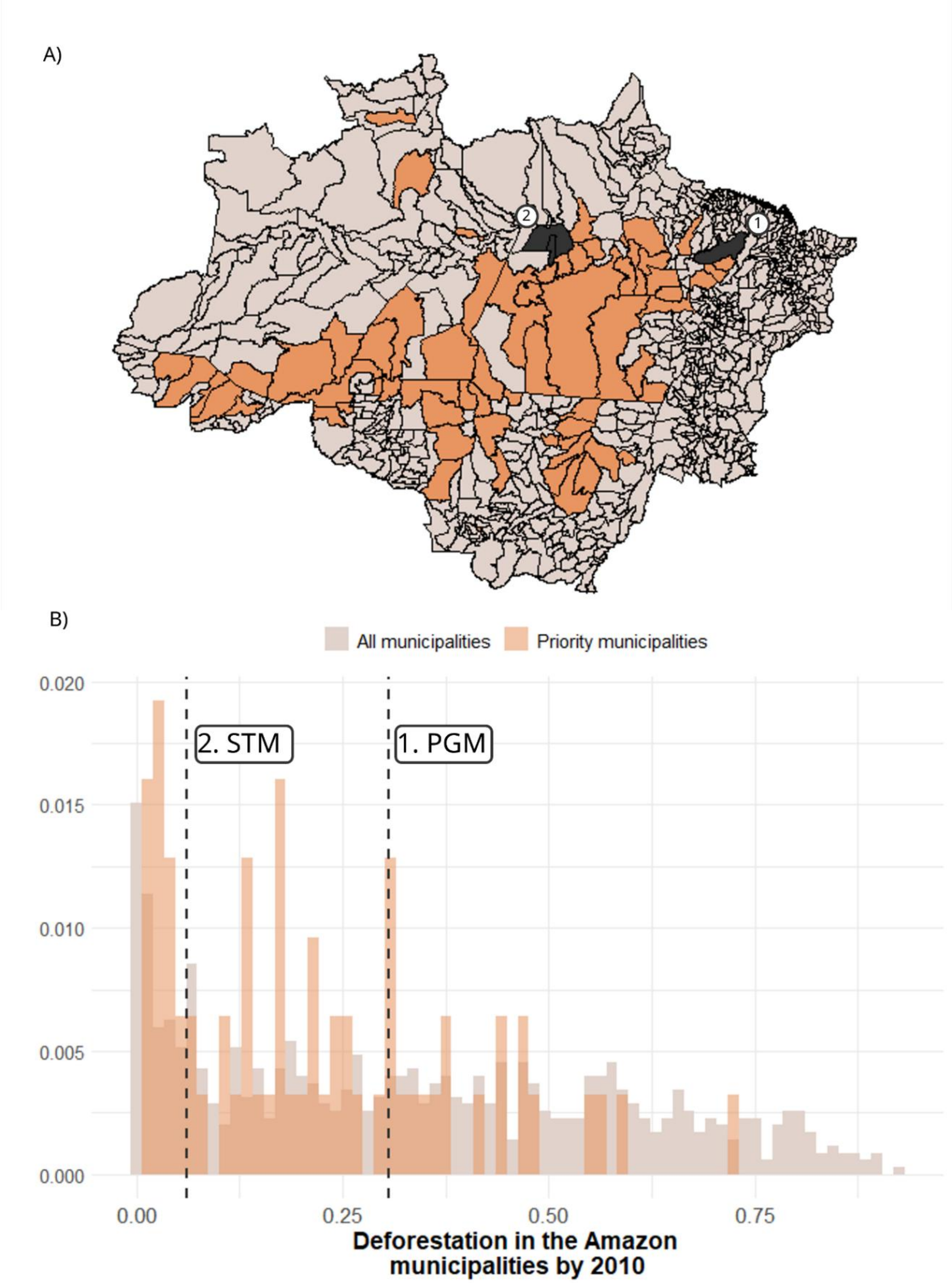
A)
2
1
B)
All municipalities
Priority municipalities
0.020
0.015
0.010
0.005
0.000
2. STM
1. PGM
0.00
0.25
0.50
0.75
Deforestation in the Amazon
municipalities by 2010

Fig. S2.

**Location of the study regions relative to deforestation patterns across municipalities in the Brazilian Amazon.** (A) Map showing all municipalities within the Brazilian Amazon region (light grey), highlighting those designated as priorities in the 'Arc of Deforestation' due to historically high deforestation rates (orange). The two study regions of this work, Paragominas (1, PGM) and Santarém (2, STM), are indicated in dark grey. Although Paragominas previously featured among priority municipalities, it has been removed from this list. (B) Histogram depicting the distribution of municipalities based on the proportion of their total area deforested by 2010. All Amazonian municipalities are shown in lighter shades, while priority municipalities are highlighted in orange. Dashed vertical lines indicate the positions of Paragominas and Santarém relative to other municipalities.

***Figure S3***

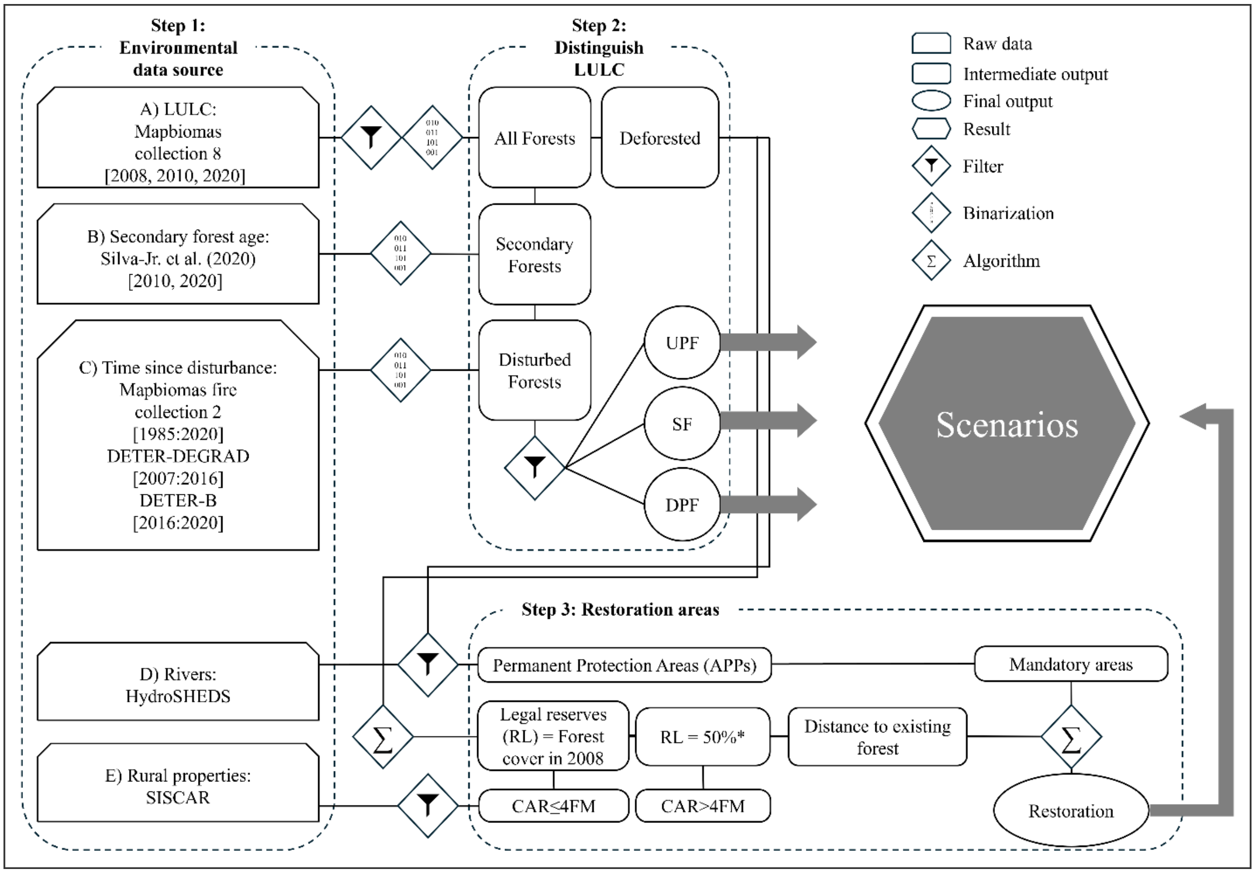


Fig. S3.

**Workflow outlining the steps involved in building the conservation scenarios.** The image summarizes the methodological approach to developing the intervention scenarios, structured into three main steps: (1) compilation of environmental data sources, including land use and land cover (LULC), secondary forest age, time since last within-forest disturbances (i.e., fire and logging), river networks, and property boundaries; (2) classification of forests types, distinguishing between secondary forests (SF), disturbed primary forests (DPF), and undisturbed primary forests (UPF); and (3) identification of potential restoration areas based on mandatory restoration targets (Permanent Protection Areas – APPs and Legal Reserves – RL) determined by law, and proximity to existing forests. The final output from these processes fed into the definition and analysis of the conservation intervention scenarios.

*Figure S4*

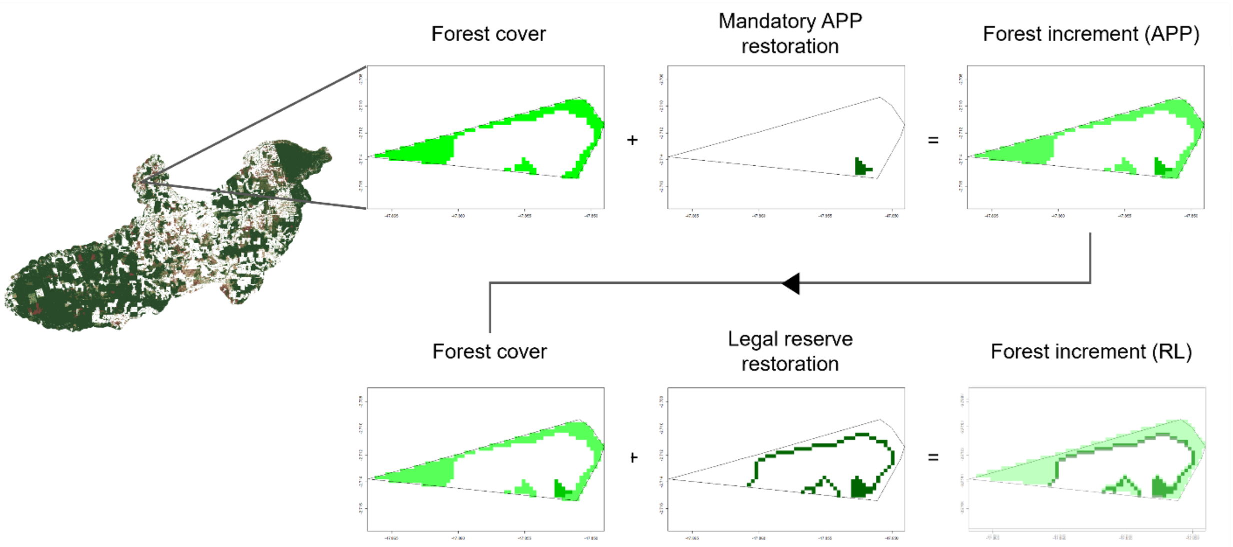


Fig. S4.

**Hierarchical steps for selecting restoration areas.** The steps include: (i) Estimating initial forest cover to identify legal reserves (RL) deficits; (ii) Adding permanent protection areas (APPs) as for mandatory restoration; (iii) Measuring the forest increment due to APP addition; (iv) Recalculating RL deficits post-APP restoration; and (v) Incrementing forest cover based on proximity to existing forests until legal compliance is achieved. Maps provide a detailed visualization at the property level in one property in Paragominas of the changes in forest cover and the strategic addition of restored areas to meet legal and ecological requirements.

*Figure S5*

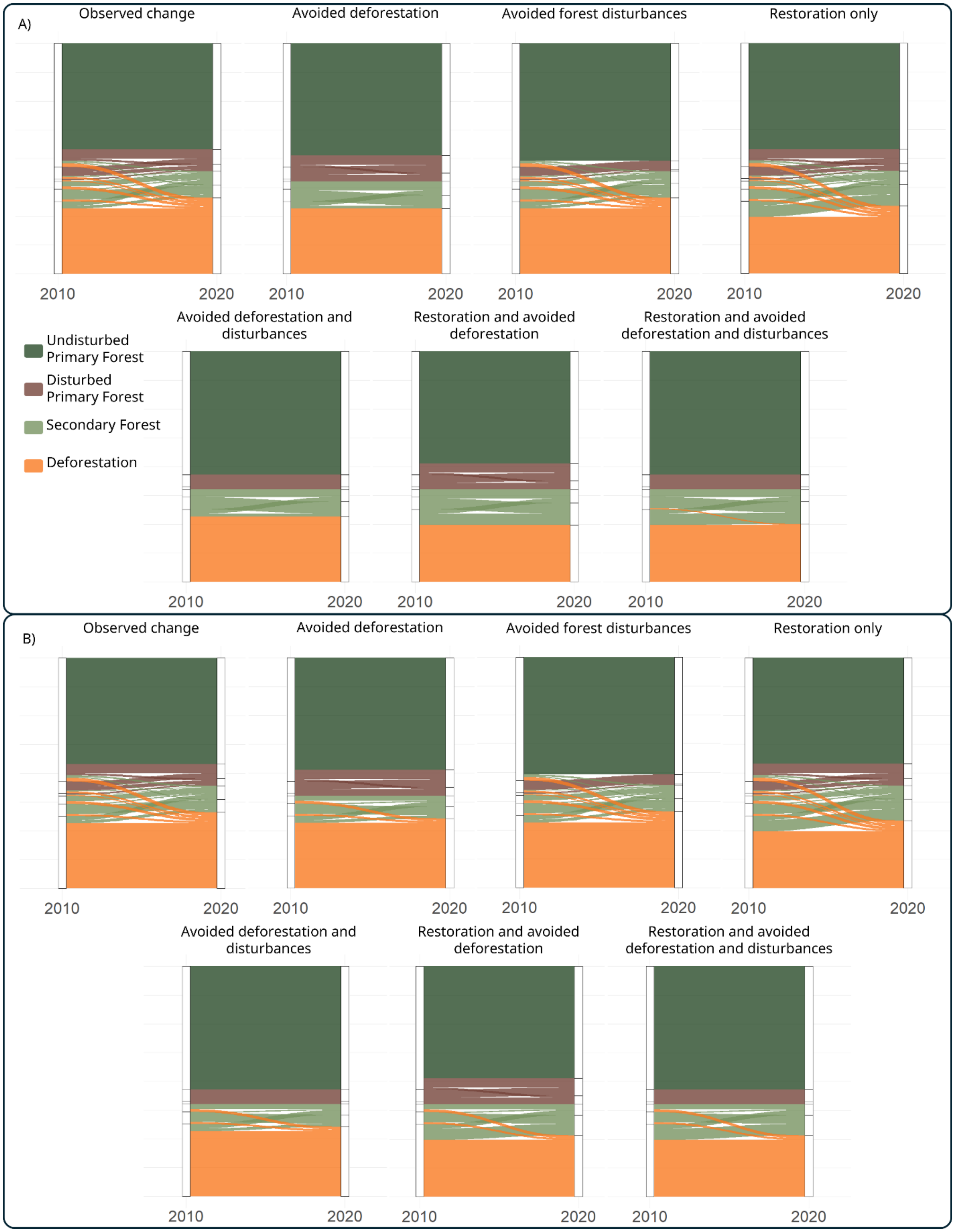

A)
Observed change
Avoided deforestation
Avoided forest disturbances
Restoration only
2010
2020
Avoided deforestation and disturbances
Restoration and avoided deforestation
Restoration and avoided deforestation and disturbances
Undisturbed Primary Forest
Disturbed Primary Forest
Secondary Forest
Deforestation
B)

Fig. S5.

**Sankey diagrams of forest type change in Paragominas from 2010 to 2020.** Each diagram depicts the transition in one of the following intervention scenarios: (i) Observed changes (i.e., what actually happen between 2010 and 2020); (ii) Avoiding deforestation; (iii) Avoiding within-forest disturbances (i.e., fire and logging); (iv) Avoiding deforestation and disturbances; (v) Restoration alone; (vi) Restoration and avoiding deforestation; and (vii) Restoration and avoiding deforestation and disturbances. (A) Scenarios considering interventions in all forest types (i.e., primary and secondary). (B) Scenarios considering interventions only in primary forest.

***Figure S6***

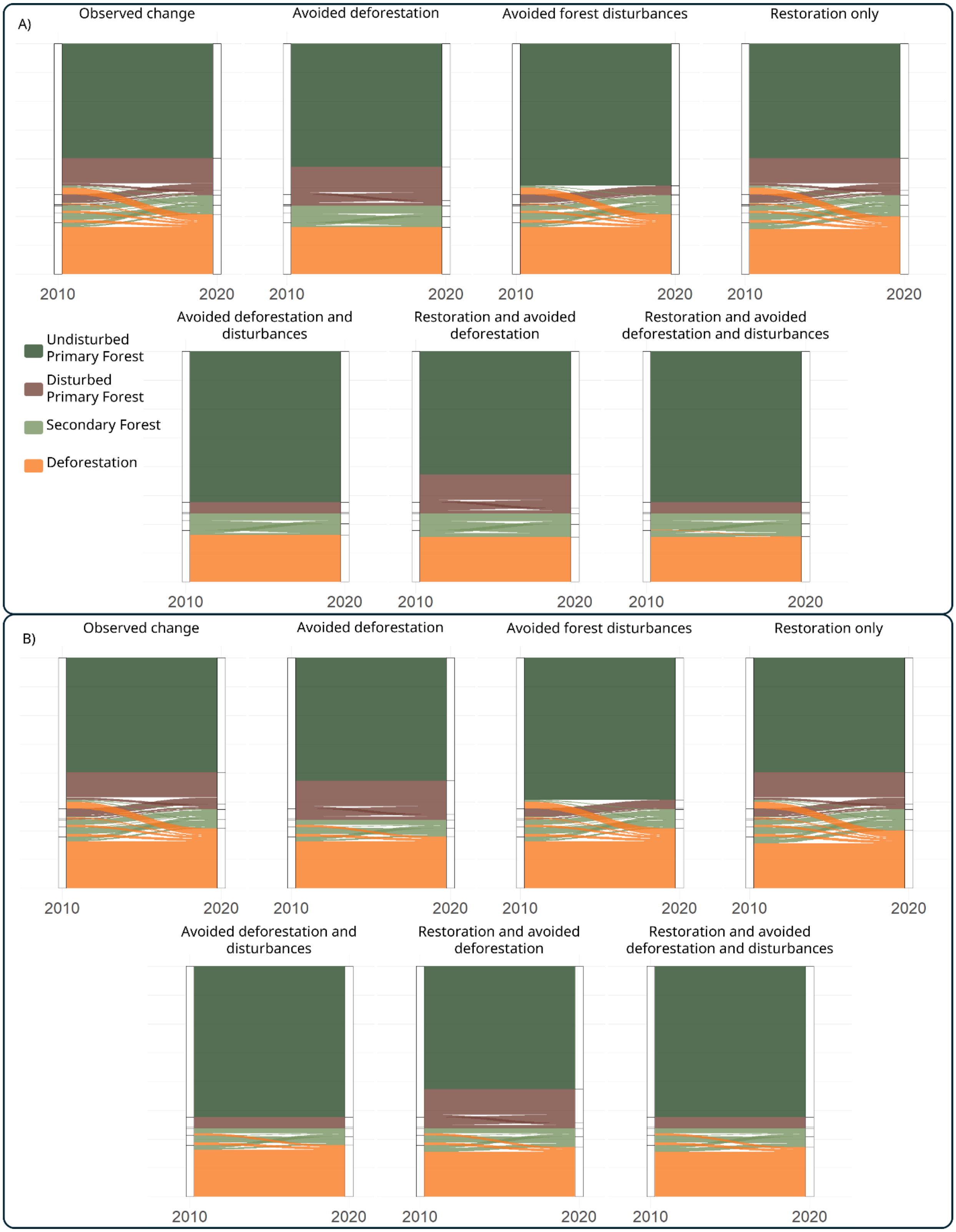
A)
Observed change
Avoided deforestation
Avoided forest disturbances
Restoration only
2010
2020
Avoided deforestation and disturbances
Restoration and avoided deforestation
Restoration and avoided deforestation and disturbances
Undisturbed Primary Forest
Disturbed Primary Forest
Secondary Forest
Deforestation
B)
Observed change
Avoided deforestation
Avoided forest disturbances
Restoration only
Avoided deforestation and disturbances
Restoration and avoided deforestation
Restoration and avoided deforestation and disturbances

Fig. S6.

**Sankey diagrams of forest type change in Santarém from 2010 to 2020.** Each diagram depicts the transition in one of the following scenarios: (i) Observed changes (i.e., what actually happen between 2010 and 2020); (ii) Avoiding deforestation; (iii) Avoiding within-forest disturbances (i.e., fire and logging); (iv) Avoiding deforestation and disturbances; (v) Restoration alone; (vi) Restoration and avoiding deforestation; and (vii) Restoration and avoiding deforestation and disturbances. (A) Scenarios considering interventions in all forest types (i.e., primary and secondary). (B) Scenarios considering interventions only in primary forest.

*Figure S7*

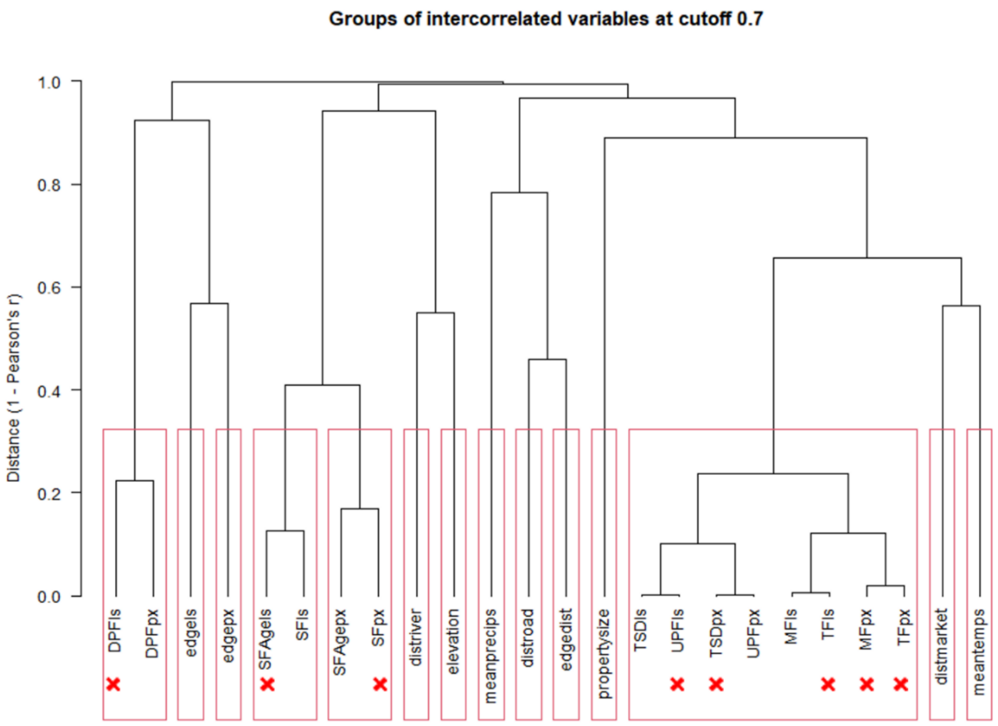


Fig. S7.

**Collinearity analysis of environmental variables.** Dendrogram and red squares represent groups of intercorrelated variables at cutoff 0.7, where the y-axis represents distances between groups (1-Pearson's r). Red crosses bellow variables represent those excluded based on variance inflation factor. Details on variables acronyms in table S1.

***Figure S8***

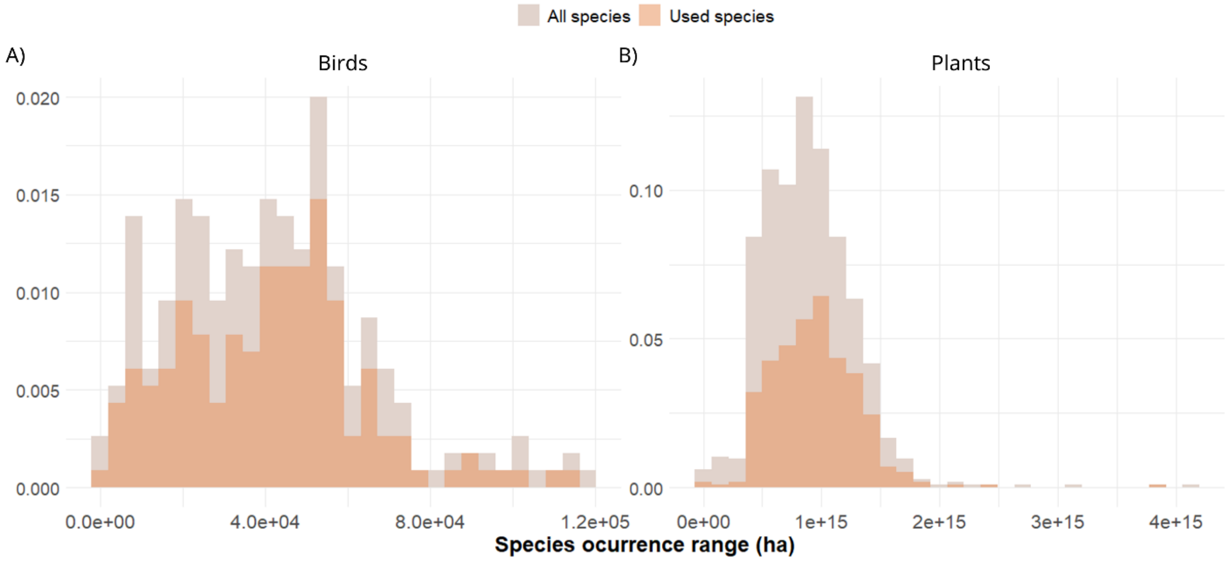


Fig. S8.

**Distribution of species occurrence ranges (ha) of forest-dependent birds and plants.** The histograms illustrate the distribution of geographical range sizes for all forest-dependent species recorded in the study regions (light shades), and the subset of species with sufficient occurrence records (orange shades) that were included in the species distribution models used for estimating biodiversity benefits. Panel A represents bird species, while panel B corresponds to plant species. Species with narrower geographical ranges were assigned greater weight in the biodiversity analysis to reflect their higher conservation importance.

***Figure S9***

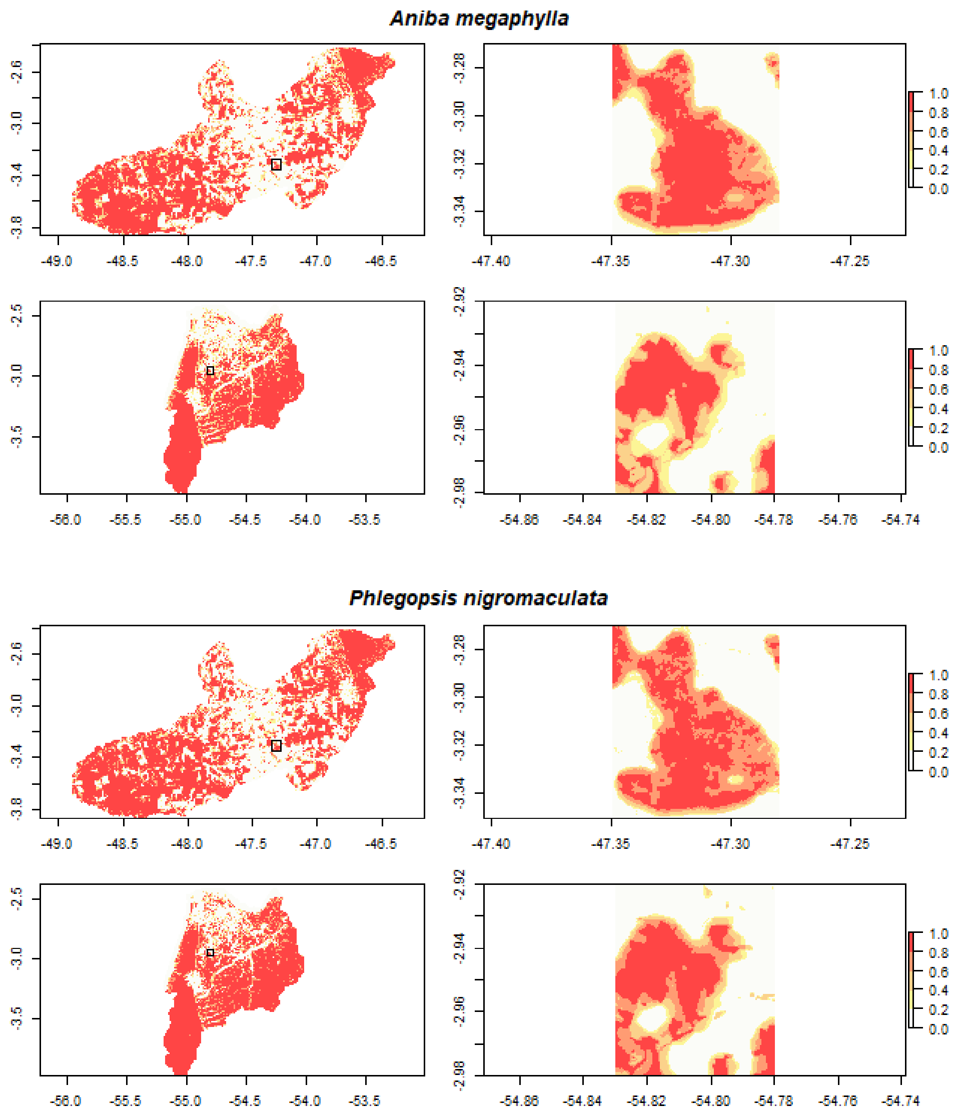


Fig. S9.

**Occurrence probabilities projections.** Maps of occurrence probabilities for two species, modelled using Random Forest algorithm. The upper panels depict occurrence probabilities for a plant species (A*niba megaphylla*) in Paragominas (i) with a zoomed-in view (ii), and in Santarém (iii) with a zoomed-in view (iv). The lower panels show habitat suitability for a bird species (*Phlegopsis nigromaculata*) in Paragominas (v) with a zoomed-in view (vi), and in Santarém (vii) with a zoomed-in view (viii).

*Figure S10*

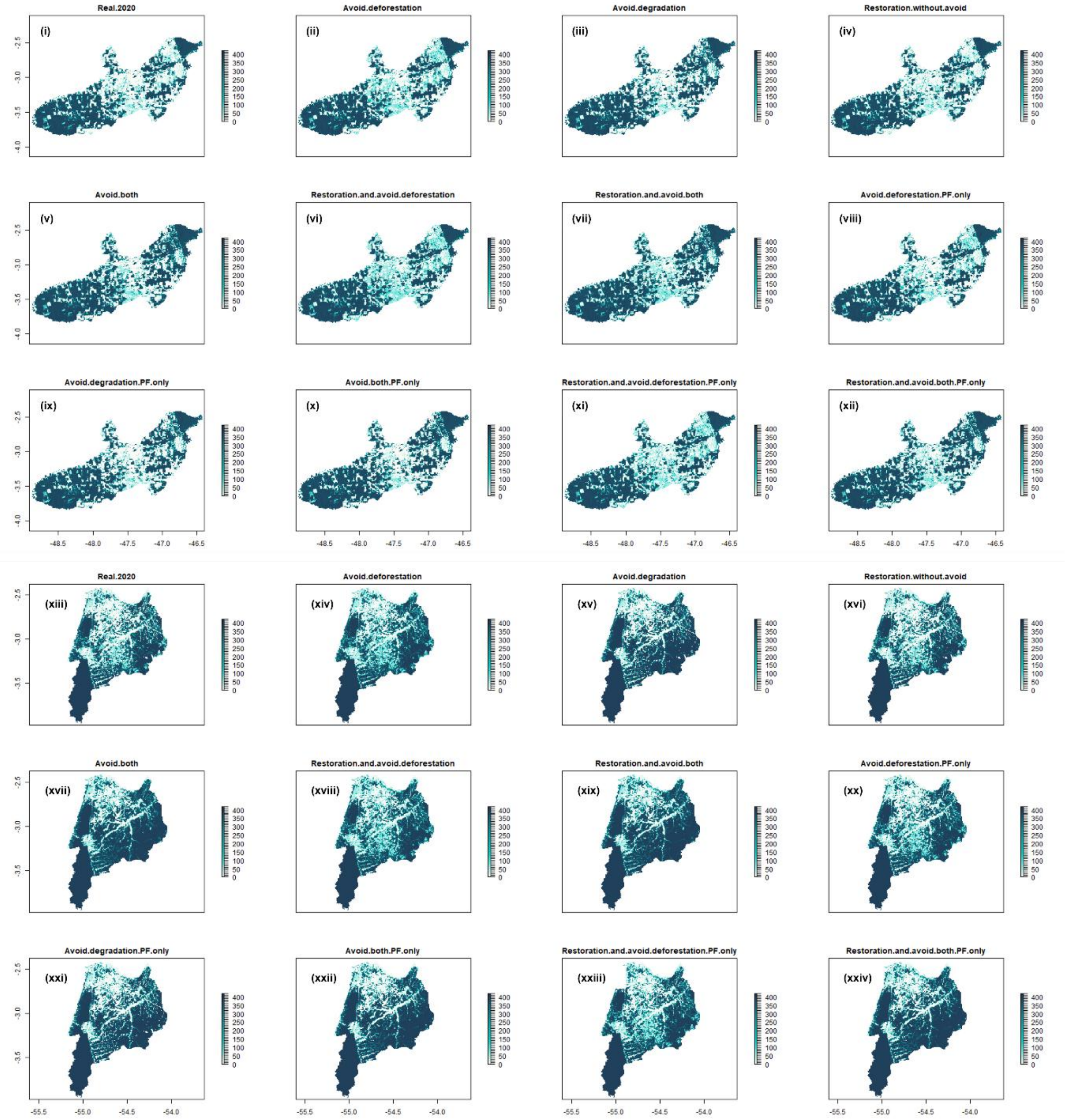


Fig. S10.

**Maps representing biodiversity conservation values.** The maps illustrate the sum of the product of habitat suitability and a scale based on inverse species' range size, across twelve different scenarios. Maps (i) to (xii) Paragominas and (xiii) to (xxiv) Santarém. The color gradient indicates areas of varying biodiversity conservation values, with darker shades representing higher values.

*Figure S11*

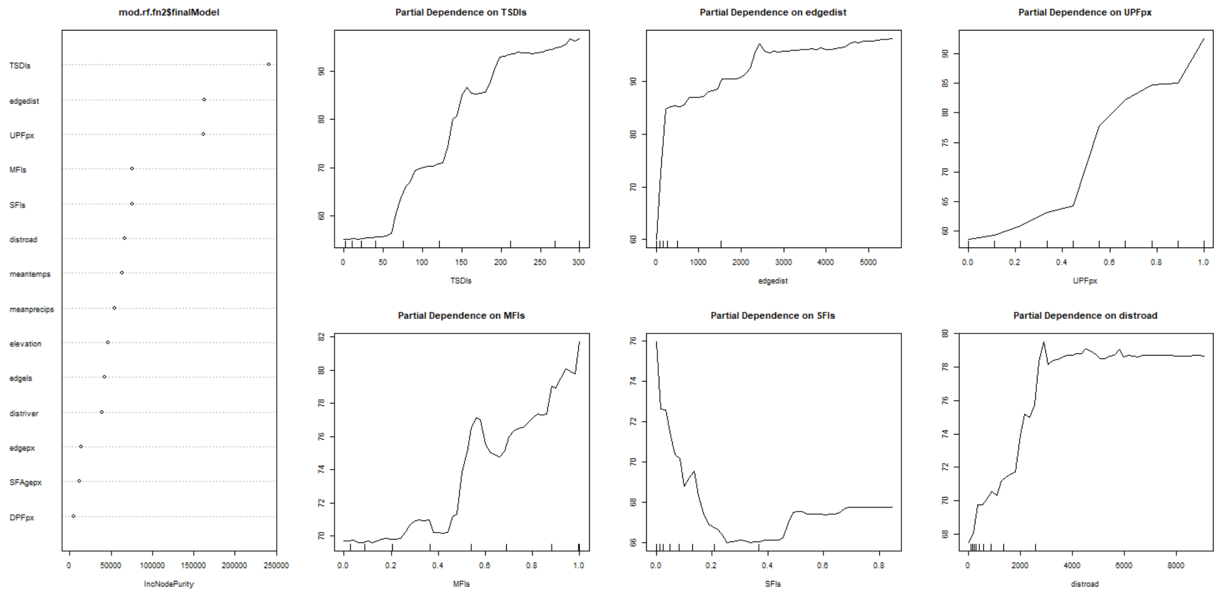


Fig. S11.

**Importance of environmental variables and partial dependence plots in the random forest model predicting above-ground carbon stocks.** Left panel displays the importance of each variable based on the mean decrease in accuracy (%IncMSE). Right panels present partial dependence plots for the six most important variables, showing their marginal effects on predicted above-ground carbon stocks. The best random forest regression model, tested with 1,700 trees and 5 variables at each split, achieved a mean of squared residuals of 1914.97 and explained 54.86% of the variance in carbon stocks.

*Figure S12*

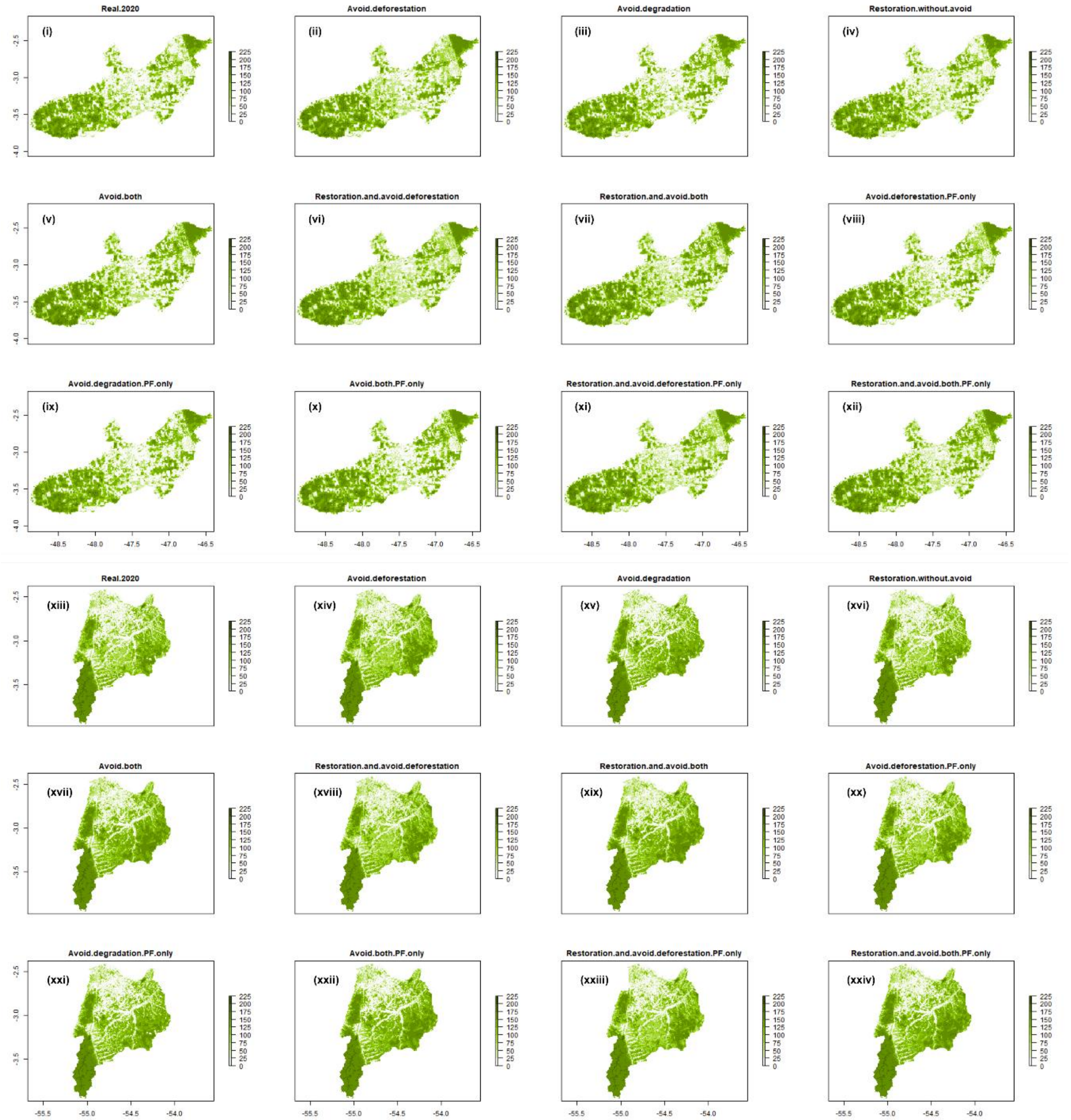


Fig. S12.

**Above-ground carbon stocks projections.** Maps illustrate the spatial distribution of predicted carbon stocks based on random forest model across twelve scenarios. Maps (i) to (xii) Paragominas and (xiii) to (xxiv) Santarém. The color gradient indicates areas of varying carbon stocks, with darker shades representing higher values.

*Figure S13*

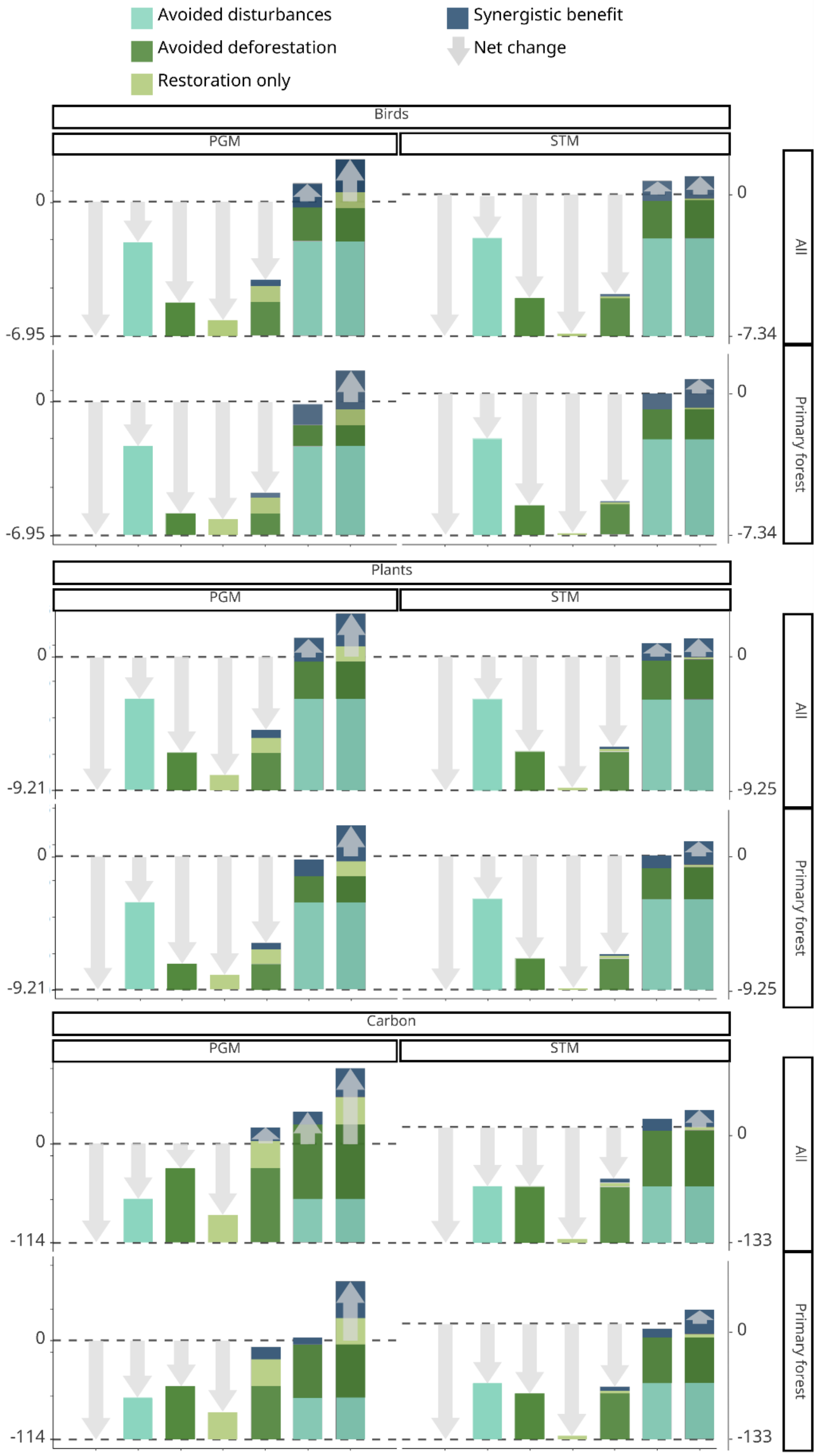
Avoided disturbances
Avoided deforestation
Restoration only
Synergistic benefit
Net change
Birds
PGM
STM
All
Primary forest
0
-6.95
-7.34
Plants
-9.21
-9.25
Carbon
-114
-133

Fig. S13.

**Summed net change and benefit in biodiversity and carbon under different conservation intervention scenarios.** Panels represent the total net change and benefits in biodiversity and carbon across the study regions of Paragominas (PGM) and Santarém (STM) over a 10-year period (2010–2020). The top panels show biodiversity net change and benefits for birds, the middle panels for plants, and the bottom panels for carbon. Each intervention was applied either across all forest types (All) or restricted to primary forests (Primary Forest). Grey arrows indicate the net change for each intervention scenario, with downward arrows denoting net losses and upward arrows indicating net gains. Avoided within-forest disturbances (light blue), avoided deforestation (green), and restoration (light green) all contribute to benefits, with the combined interventions generating synergistic gains (dark blue), which exceed the sum of individual interventions. Dashed lines represent the 2010 baseline (upper line) and the observed changes from 2010 to 2020 (lower line).

*Figure S14*

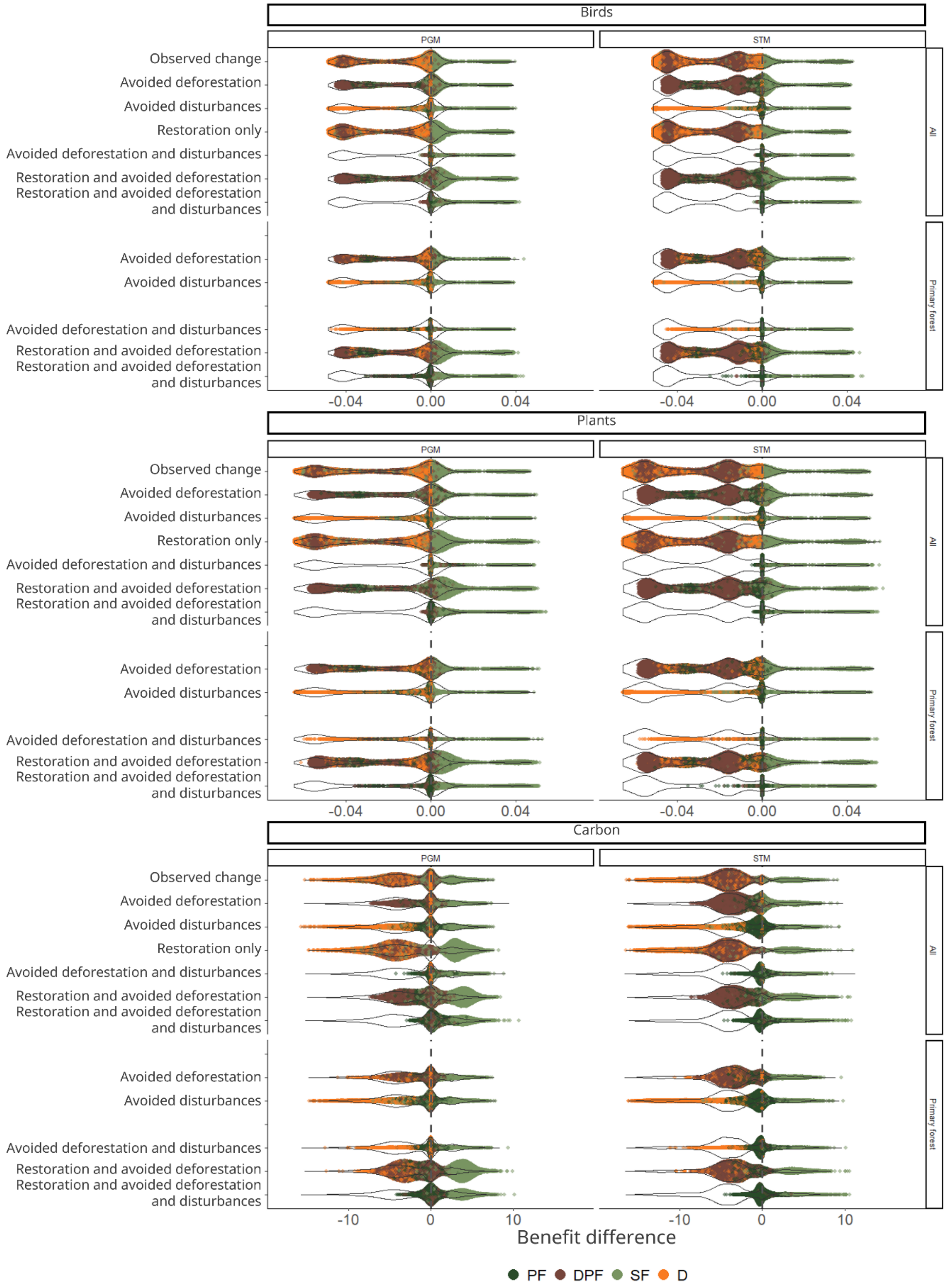

Birds
Plants
Carbon
PGM
STM
All
Primary forest
Observed change
Avoided deforestation
Avoided disturbances
Restoration only
Avoided deforestation and disturbances
Restoration and avoided deforestation
Restoration and avoided deforestation and disturbances
-0.04
0.00
0.04
-10
0
10
Benefit difference
PF
DPF
SF
D

Fig. S14.

**Pixel-level distribution of net changes in biodiversity and carbon benefits across intervention scenarios.** Plots illustrate the pixel-level distribution of net changes in biodiversity and carbon under different conservation intervention scenarios for Paragominas (PGM) and Santarém (STM). The upper panels show biodiversity net change for birds, the middle panels for plants, and the lower panels for carbon. Each intervention was applied either across all forest types (All) or only to primary forests (Primary Forest). The y-axis displays the interventions, while the x-axis represents the net change relative to the baseline. The outlined violin plot is identical across all interventions, representing the distribution of values for the predicted net change between 2010 and 2020. Inside each violin plot, dots represent individual pixels that changed their forest type during this period. Dots are colored according to forest type: undisturbed primary forest (UPF), disturbed primary forest (DPF), secondary forest (SF), and deforested areas (D). In the predicted 2010-2020 net change, the dots naturally fit the shape of the violin plot, aligning with the predicted net change distribution. However, in the other intervention scenarios, dots were constrained within the violin plot to highlight the net change difference between the predicted 2010-2020 net change and the intervention scenarios. The maximum width of dot dispersal had to conform to the shape of the violin plot and high pixel densities around zero (i.e., areas of little to no net loss) resulted in a visual difference of positive contributions. The vertical dashed line represents the 2010 baseline, with values below the baseline indicating net loss and values above representing net gain.

*Figure S15*

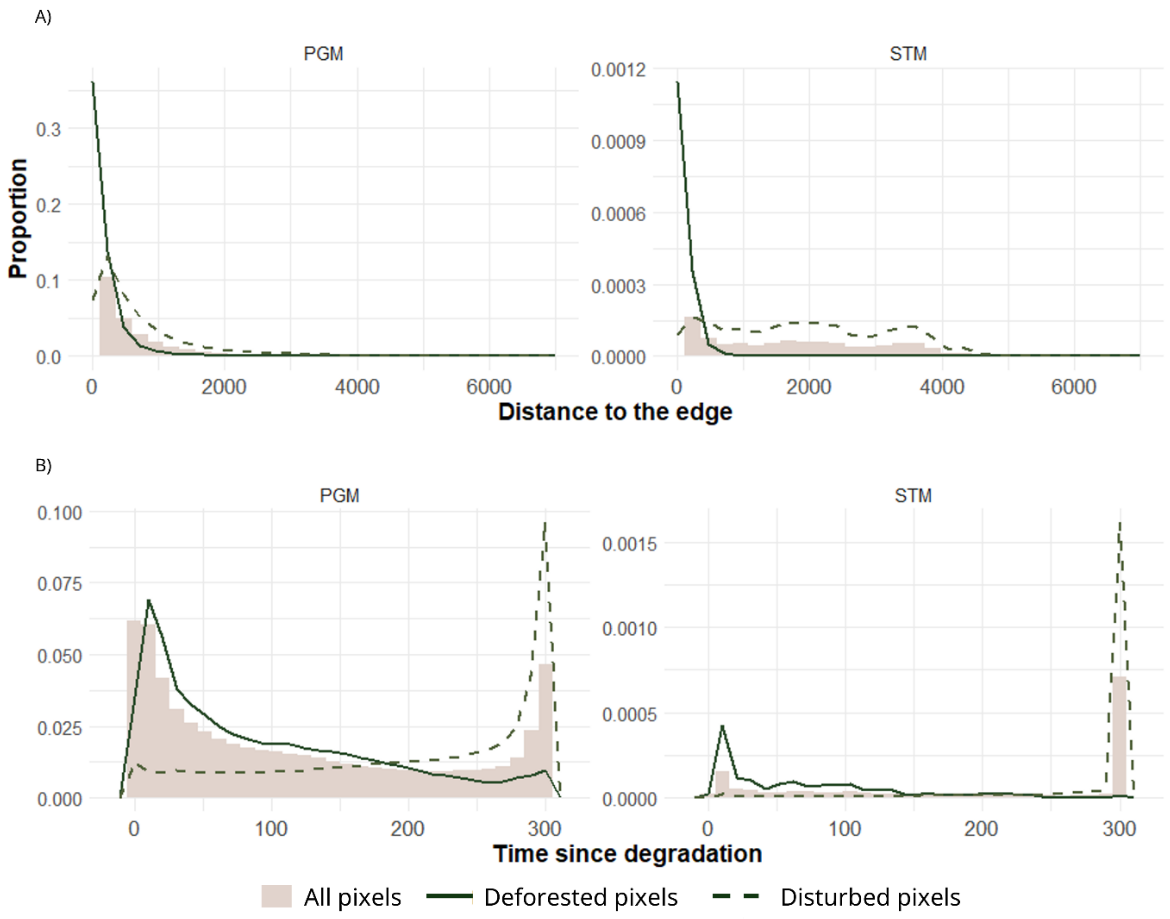


Fig. S15.

**Spatial and temporal characteristics of deforestation and disturbance events in the study regions.** The histograms illustrate the distribution of (A) distance to forest edges (m) and (B) time since the last within-forest disturbance event (years) across all pixels in Paragominas (PGM, left) and Santarém (STM, right). Light grey bars represent the distribution for all pixels, while solid lines indicate the characteristics of pixels where deforestation occurred between 2010 and 2020, and dashed lines represent pixels that experienced disturbance. In both regions, deforestation was highly concentrated near forest edges, with most events occurring within 500 m from the edge and in areas with a short time since the last disturbance. In contrast, disturbance events occurred relatively farther from the forest edge, particularly in Santarém, and often in areas with no prior recorded disturbance.

***Figure S16***

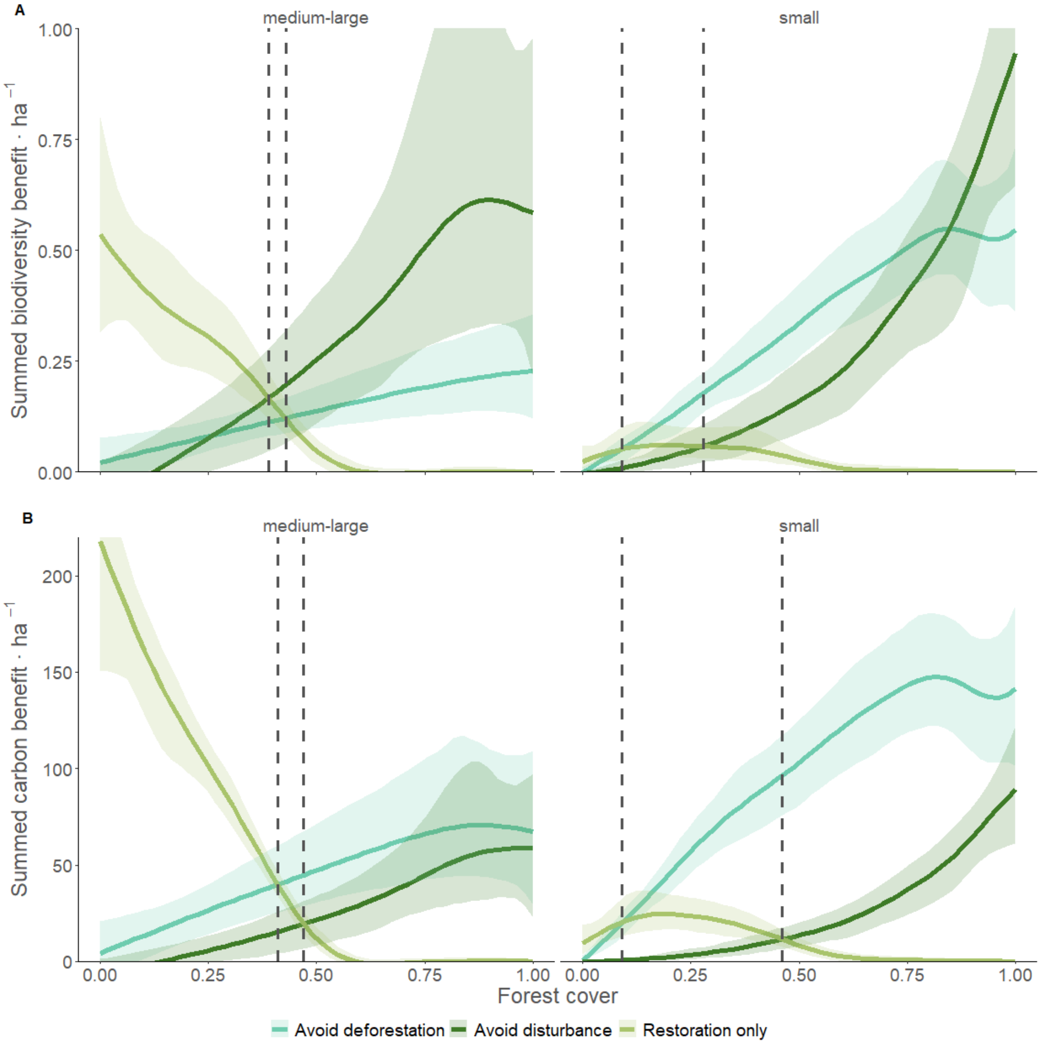


Fig. S16.

**Variation in the summed benefit per-hectare of property of each intervention as a function of property size and forest cover.** (A) Biodiversity and (B) carbon benefits for each intervention scenario (avoid deforestation, avoid within-forest disturbance, and restoration only). Panels on top show observed bootstrapped means and 95% confidence intervals across 1,000 iterations; panels below show null model results based on randomized region, size, and forest cover. Vertical dashed lines indicate the crossover points where restoration begins to outperform avoided deforestation or avoided disturbances.

***Figure S17***

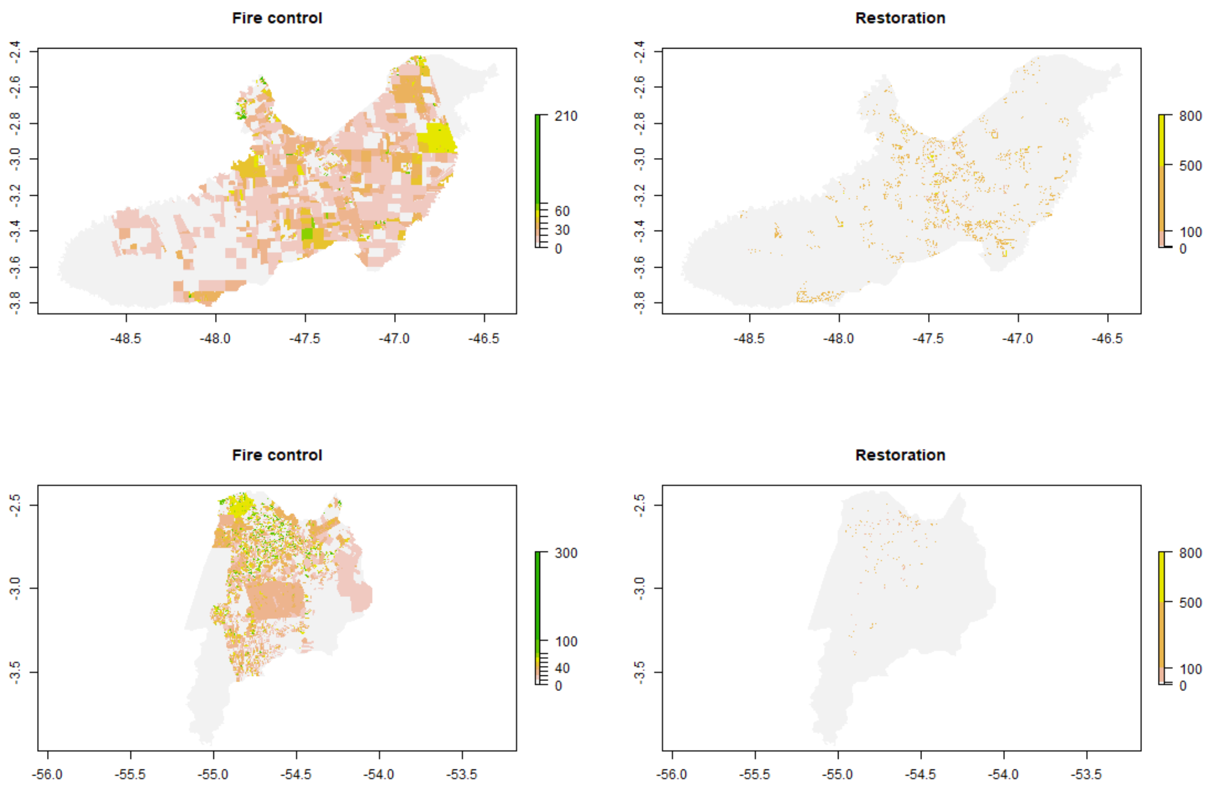


Fig. S17.

**Management costs for fire control and restoration.** The maps depict the spatial distribution of implementation costs associated with fire control (left) and restoration (right) for Paragominas (top) and Santarém (bottom). These costs are represented by a color gradient, with higher costs shown in green and yellow.

***Figure S18***

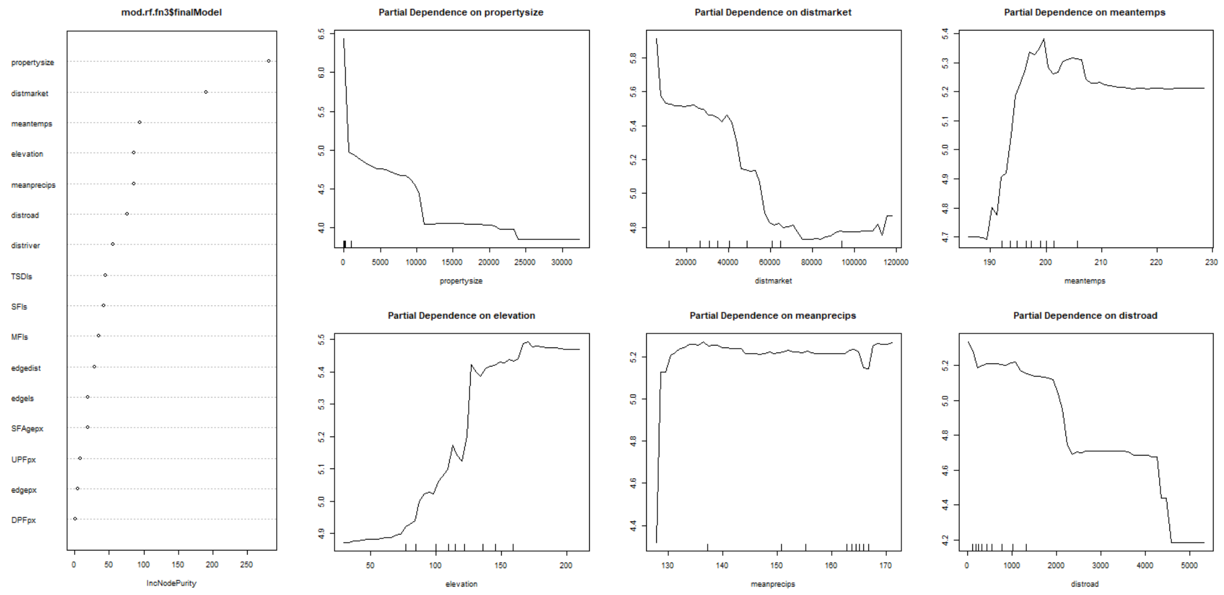


Fig. S18.

**Importance of environmental variables and partial dependence plots in the random forest model predicting foregone farm income.** Left panel displays the importance of each variable based on the mean decrease in accuracy (%IncMSE). Right panels present partial dependence plots for the six most important variables, showing their marginal effects on predicted foregone farm income. The best random forest regression model, tested with 1,750 trees and 7 variables at each split, achieved a mean of squared residuals of 2,336.76 and explained 18.42% of the variance in foregone farm income.

*Figure S19*

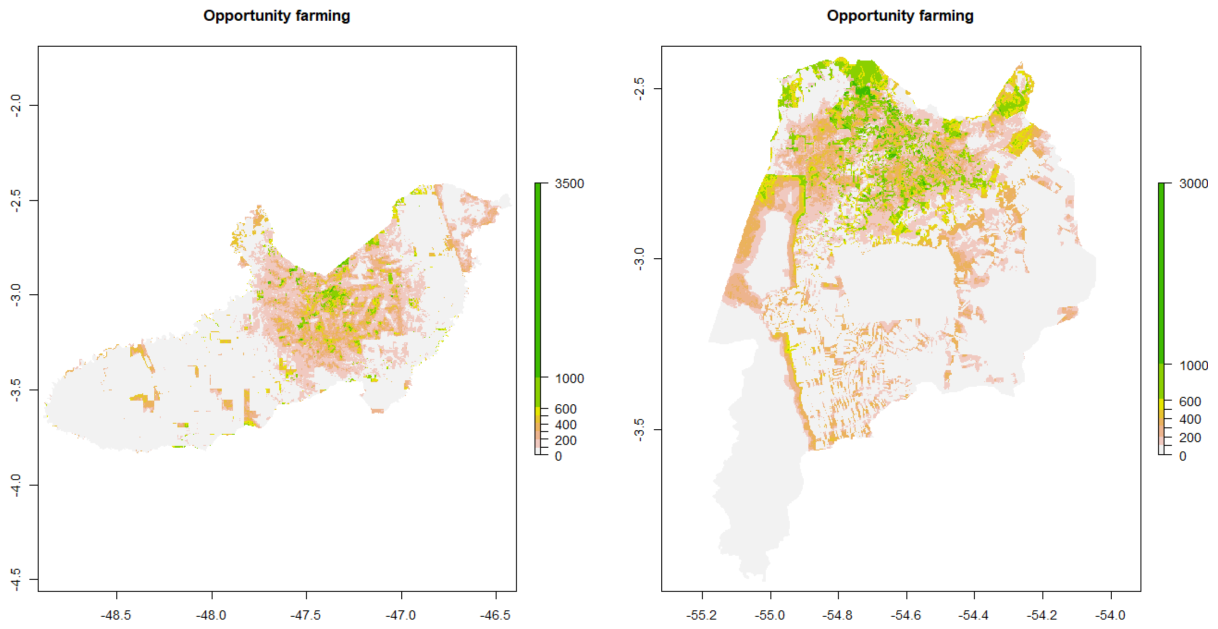


Fig. S19.

**Opportunity costs linked to foregone farm income projections.** The maps depict the spatial distribution of opportunity costs associated with foregone farm income, based on the predictions from the random forest model for Paragominas (left) and Santarém (right). The color gradient represents varying levels of economic impact, with higher opportunity costs shown in green.

*Figure S20*

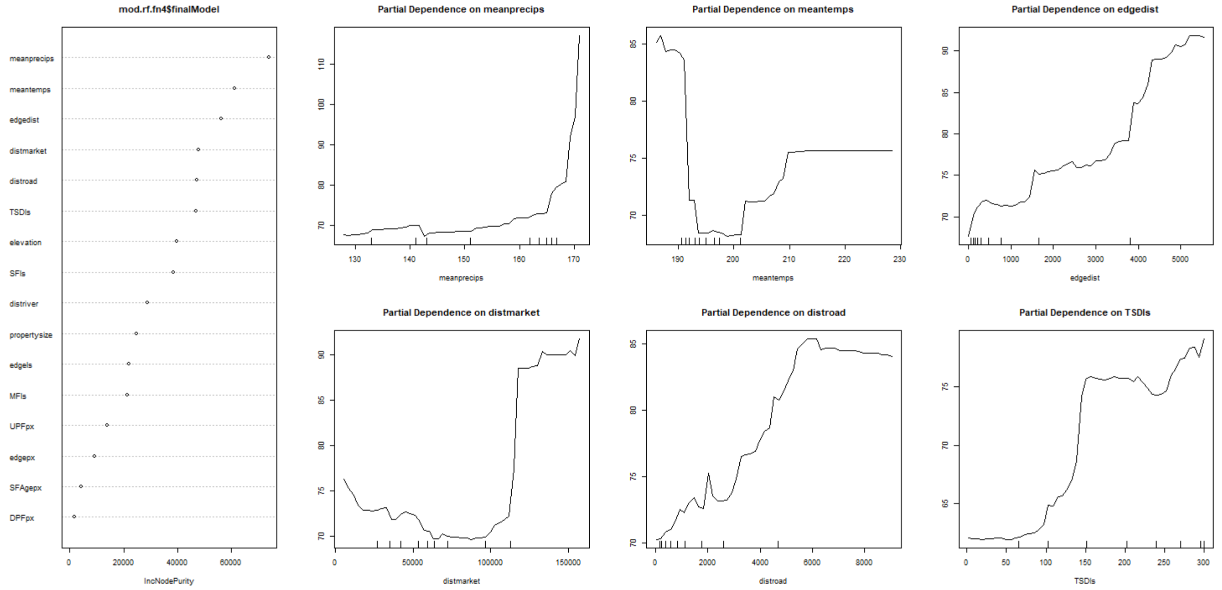


Fig. S20.

**Importance of environmental variables and partial dependence plots in the random forest model predicting foregone logging income.** Left panel displays the importance of each variable based on the mean decrease in accuracy (%IncMSE). Right panels present partial dependence plots for the six most important variables, showing their marginal effects on predicted foregone logging income. The best random forest regression model, tested with 1,350 trees and 3 variables at each split, achieved a mean of squared residuals of 2,957.63 and explained 11.87% of the variance in foregone logging income.

*Figure S21*

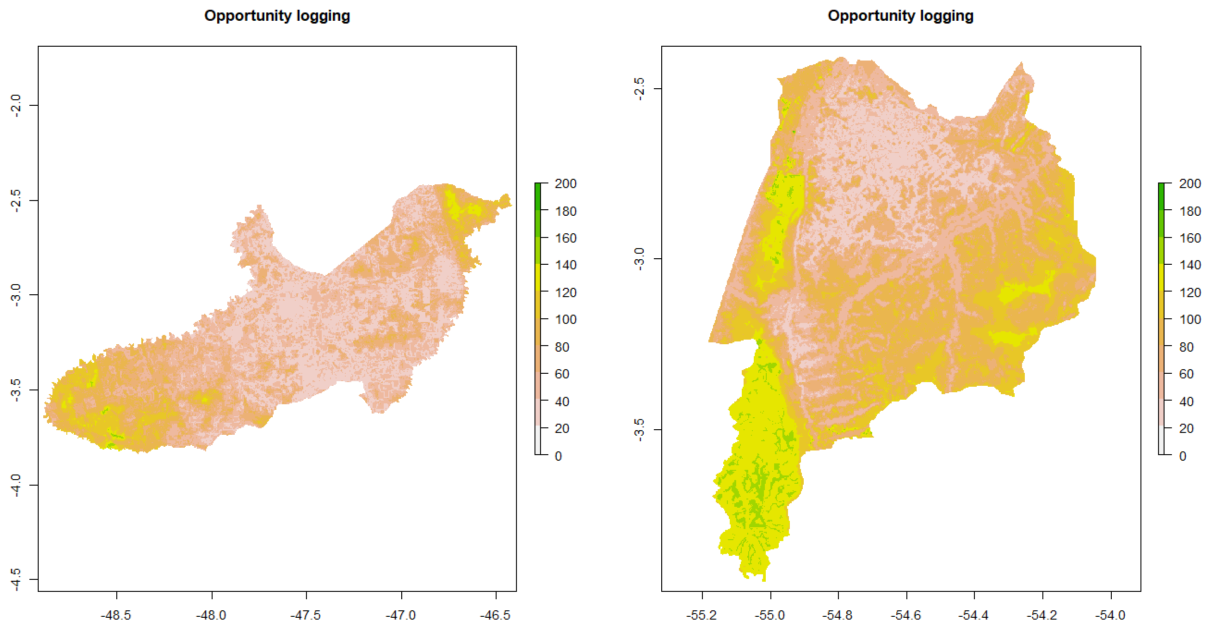


Fig. S21.

**Opportunity costs linked to foregone logging income projections.** The maps depict the spatial distribution of opportunity costs associated with foregone logging income, based on the predictions from the random forest model for Paragominas (left) and Santarém (right). The color gradient represents varying levels of economic impact, with higher opportunity costs shown in green.

*Figure S22*

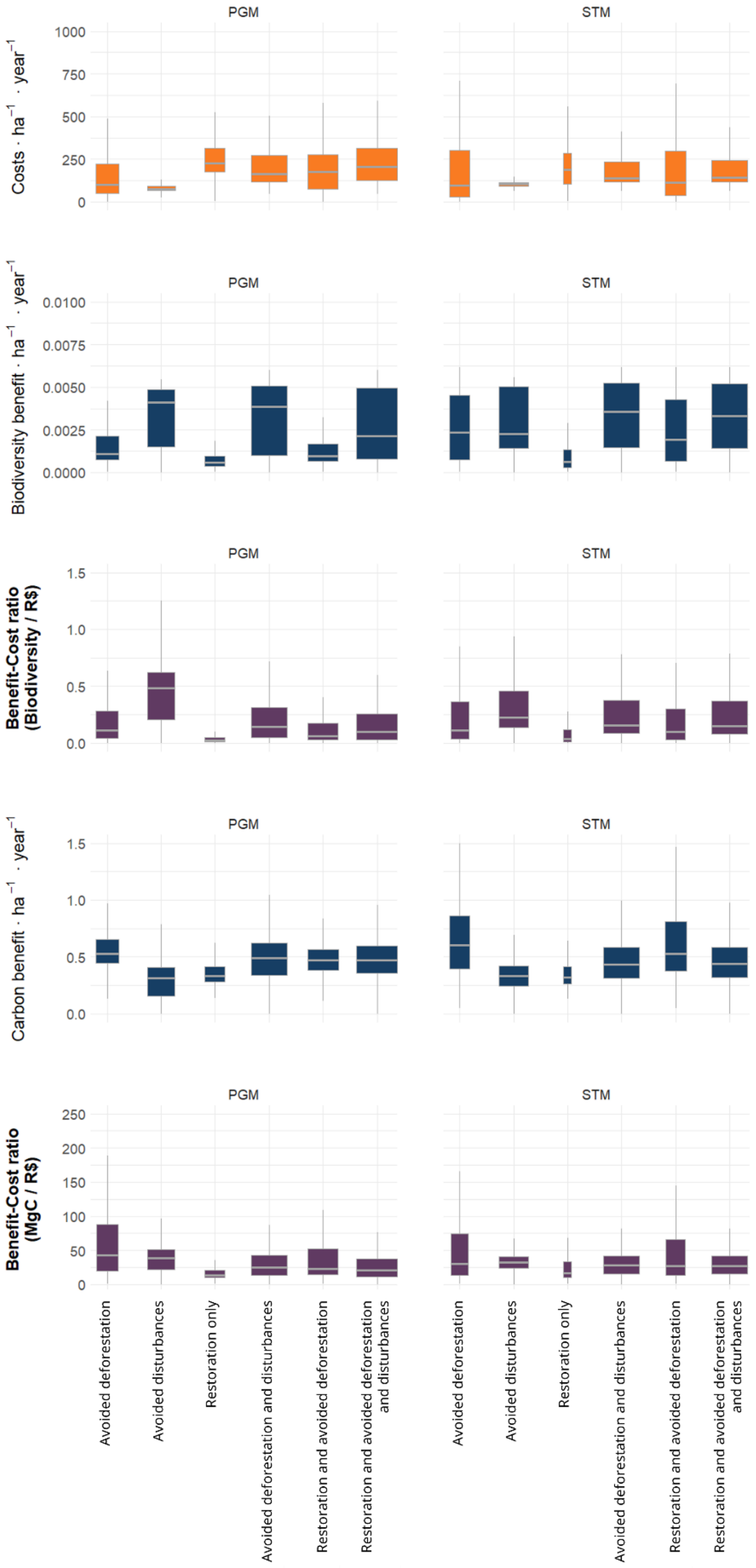

PGM
STM
Costs · ha⁻¹ · year⁻¹
1000
750
500
250
0
Biodiversity benefit · ha⁻¹ · year⁻¹
0.0100
0.0075
0.0050
0.0025
0.0000
Benefit-Cost ratio (Biodiversity / R$)
1.5
1.0
0.5
0.0
Carbon benefit · ha⁻¹ · year⁻¹
Benefit-Cost ratio (MgC / R$)
200
150
100
50
Avoided deforestation
Avoided disturbances
Restoration only
Avoided deforestation and disturbances
Restoration and avoided deforestation
Restoration and avoided deforestation and disturbances

Fig. S22.

**Comparative costs, per-area benefits, and cost-benefit ratios across intervention scenarios.** From top to bottom, boxplots show: (1) estimated costs per hectare per year (R$), (2) biodiversity benefits per hectare per year, (3) biodiversity benefit–cost ratio (forest biodiversity per R$10,000.00), (4) carbon benefits per hectare per year, and (5) carbon benefit–cost ratio (Mg C per R$10,000.00). Left-hand panels correspond to Paragominas (PGM) and right-hand panels to Santarém (STM).

**Figure S23**

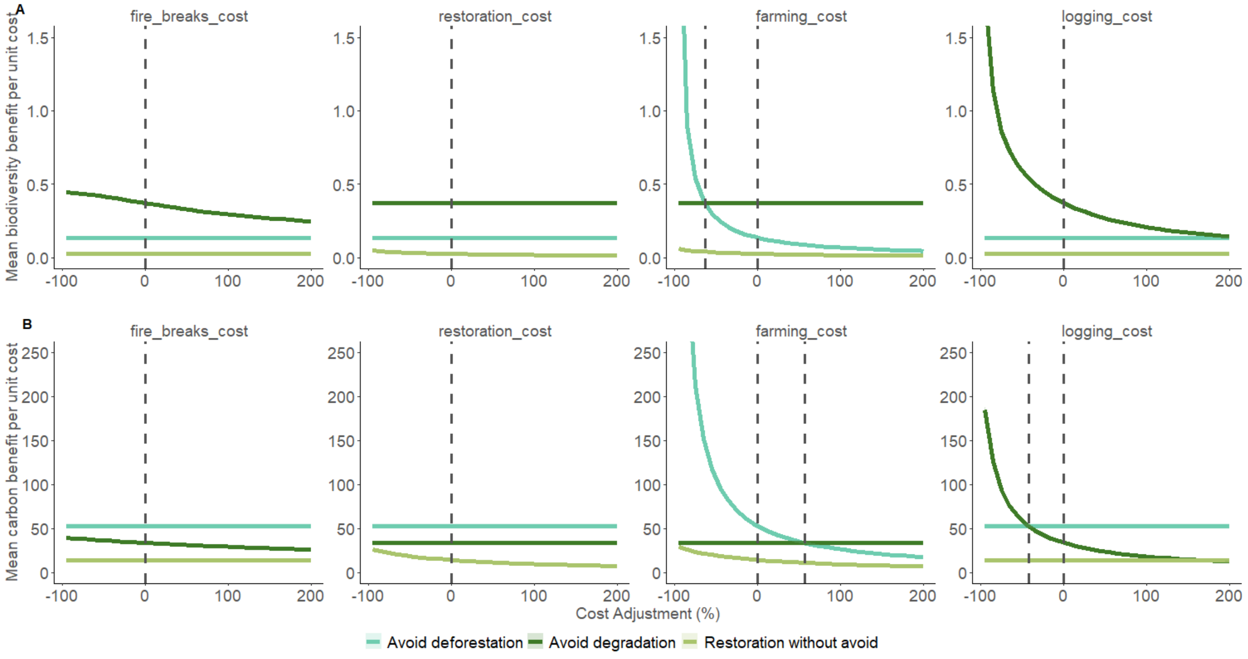


Fig. S23.

**Figure S22. Sensitivity of intervention scenario benefit-cost ratio to variation in economic assumptions for (A) biodiversity and (B) carbon benefits.** Each panel shows the mean benefit per unit cost (y-axis) across a range of percentage adjustments (x-axis) applied to one of four cost types: implementation costs of fire breaks and restoration, and opportunity costs of farming and logging. For each cost component, we modified baseline costs from –95% to +200% in 10% increments, running 100 bootstrap iterations per adjustment across 100 random subsamples of 200,000 pixels each. Lines represent the average benefit-cost ratio across all iterations, and vertical dashed lines indicate the unadjusted (baseline) cost assumption and the crossover points where shifts in the cost component alters the benefit-cost ratio between interventions.

#### 3. Supplementary Tables

##### Table S1

The predictor variables used to fit random forest models and their sources.

**Table S1**

| Predictor variable | Details | Sources |
|---|---|---|
| **UPFpx** | **Proportion of old growth primary forest (pixel scale[a]).** | **Mapbiomas derived** |
| UPFls | Proportion of old growth primary forest (landscape scale[b]) | Mapbiomas derived |
| **DPFpx** | **Proportion of disturbed primary forest (pixel)** | **Mapbiomas fire, DEGRAD; DETER-B** |
| DPFls | Proportion of disturbed primary forest (landscape) | Mapbiomas fire, DEGRAD; DETER-B |
| TSDpx | Mean time since last disturbance event (pixel) | Mapbiomas fire, DEGRAD; DETER-B |
| **TSDls** | **Mean time since last disturbance event (landscape)** | **Mapbiomas fire, DEGRAD; DETER-B** |
| SFpx | Proportion of secondary forest (pixel) | Silva Jr. et al. (2020) |
| **SFls** | **Proportion of secondary forest (landscape)** | **Silva Jr. et al. (2020)** |
| **SFagepx** | **Mean age of secondary forest (pixel)** | **Silva Jr. et al. (2020)** |
| SFagels | Mean age of secondary forest (landscape) | Silva Jr. et al. (2020) |
| TFpx | Density of total forest [= UPF + DPF + SF > 2 years old] (pixel) | |
| TFls | Density of total forest [= UPF + DPF + SF > 2 years old] (landscape) | |
| MFpx | Density of mature forest [= UPF + DPF + SF > 10 years old] (pixel) | |
| **MFls** | **Density of mature forest [= UPF + DPF + SF > 10 years old] (landscpe)** | |
| **edgepx** | **Density of forest edges base on mature forest (pixel)** | |

| | | |
|---|---|---|
| **edgels** | **Density of forest edges base on mature forest (landscape)** | |
| **edgedist** | **Mean distance to edge** | |
| **distriver** | **Mean distance to water** | **HydroSHEDS derived** |
| **distroad** | **Mean distance to road** | **Imazon[c]** |
| **elev** | **Digital elevation model (SRTM)** | **U.S. Geological Survey** |
| **meantemps** | **Mean land surface temperature** | **Nasa Earth Observation** |
| **meanprecips** | **Mean rainfall** | **Nasa Earth Observation** |
| **distmarket[d]** | **Distance to urban centre** | **IBGE** |
| **propertysize[d]** | **Property size** | **SISCAR derived** |

a. 3x3 pixel neighbourhood (~100 m buffer);
b. 35x35 pixel neighbourhood (~1 km buffer);
c. contact Imazon for access;
d. predictors used only in foregone incomes;
variables in bold selected after testing for collinearity.

#### Table S2

Summary of data and species distribution model performance statistics. ID: unique number for each species; Binomial: species scientific name without space; Group: Birds or Trees; Nrec: number of occurrences used to fit the model; *mtry* and *ntree*: optimized random forest parameters; TP: true presences; FP: false presences; TN: true absences; FN: false absences; ROC: model performance index; distriver-UPFpx: variable importance (details on variables acronyms in table S1); method: whether or not species were used in the SDMs; Shape_Area: area in km of the species occurrence range; Shape_Area_scaled: scale applied to the species occurrence range, from 0.999 to 0.001.
